# In vitro–reconstituted Drosophila Arc capsids deliver gene editors to dystrophic muscle

**DOI:** 10.64898/2026.05.25.727180

**Authors:** Blake Lash, Daniel Strebinger, Michael Segel, Samuel Chau-Duy-Tam Vo, Katherine DeLong, Julie Pham, Charles Swan, Pradeep Kumar, Yugang Zhang, Catherine C. Liu, Joanie Mok, Rhiannon Macrae, Feng Zhang

## Abstract

The intracellular delivery of therapeutic macromolecules remains a major challenge in biomedicine. Here we reconstitute the *Drosophila melanogaster* Arc1 (dArc1) retroelement-derived capsid entirely from purified recombinant protein components and in vitro transcribed RNA, creating a fully defined, cell-free assembled protein nanoparticle system. Through affinity engineering of the dArc1 RNA-binding domain, we enable efficient encapsulation of mRNA payloads and Cas9 ribonucleoproteins. Unexpectedly, we discover that dArc1 capsids bind mammalian cells through a direct interaction with the surface receptor SORCS2. Leveraging this interaction, we show that intramuscular injection of dArc1 capsids carrying Cas9 gene editors achieves up to 18% exon skipping and restores dystrophin expression in muscle fibers of mdx mice, a model of Duchenne muscular dystrophy. Enhanced delivery efficiency in regenerating and dystrophic muscle correlates with upregulated SORCS2 expression, supporting our finding that SORCS2 facilitates cellular uptake of dArc1 capsids. This work demonstrates the potential of in vitro-assembled protein nanoparticles for delivery of molecular cargoes.

## Introduction

Transposable elements make up a large proportion of eukaryotic genomes, and in some cases, hosts have co-opted these genes for important physiological functions. Among these, several retroelement-derived genes retain the capacity to encode proteins that self-assemble into virus-like capsids. The *Drosophila* Arc1 (dArc1) gene, derived from a Ty3 retrotransposon, encodes one such protein, which has been shown to self-assemble into 240-mer T=4 icosahedral capsids that package and transfer dArc1 mRNA between cells at the neuromuscular junction (NMJ)^1,2^. In its native context, capsid-mediated transfer of dArc1 mRNA is essential for synaptic plasticity at the NMJ. This activity closely resembles that of other natural systems for the transfer of nucleic acids, including viruses, raising the possibility that domesticated retroelement capsids may be adapted and engineered for delivery of genetic material.

Previous studies have made progress toward this goal, including work engineering systems based on mammalian Arc and PEG10^3,4^. However, like viral vectors, these systems rely on cell-based production in which cargo is packaged during expression in mammalian cells, posing significant challenges for manufacturing scalability ^5^. The resulting particles also lack intrinsic tropism for target cells, requiring cationic peptides or pseudotyping with viral fusogens for cellular uptake ^3,6^. dArc1 possesses a distinctive property that may circumvent this limitation: its capsid protein can be recombinantly expressed in *E. coli* and reconstituted into virus-like particles in vitro ^2^, raising the possibility of a fully cell-free, scalable assembly pipeline for nucleic acid delivery.

Here, we describe the development of in vitro–assembled protein nanoparticles (PNPs) based on an engineered dArc1 (henceforth dArc) protein. We engineer dArc capsids for efficient RNA and ribonucleoprotein (RNP) encapsulation and discover that these capsids natively bind and deliver cargo to mammalian cells through the surface receptor SORCS2. We show that dArc capsids can deliver Cas9 RNPs to muscle cells, and harness this endogenous binding to deliver genetic medicines to muscle *in vivo*.

### dArc self-assembles *in vitro* into capsids, which can be engineered for mRNA and RNP delivery

To develop a cell-free assembly pipeline, we purified recombinant dArc protein from *E. coli* fused to a maltose-binding protein (MBP) and an engineered SUMO tag (bdSUMO) tag ^7^ to increase solubility and prevent premature self-assembly during production. We observed capsid formation upon removal of the tag with engineered SUMO protease (bdSENP), in line with previous work ^2^ (Fig. 1, a and b). Although the native capsid binds and packages mRNA naturally, it is unknown whether RNA is required to template assembly of the capsid *in vitro*. We tested whether exogenous IVT *arc1* mRNA (+/− mRNA) and a polyethyleneimine (PEI) precipitation step to strip contaminating *E. coli* nucleic acids during purification (+/− PEI) affected assembly (Fig. 1c, Supplementary Fig. 1b). Without exogenous mRNA, capsids formed only when the PEI step was omitted (Fig. 1c), suggesting that co-purifying bacterial nucleic acids sufficed to template assembly. With PEI precipitation (Supplementary Fig. 1a), little capsid formation occurred upon tag cleavage, but this was rescued by adding IVT arc1 mRNA — indicating that mRNA is required for dArc capsid assembly (Fig. 1c, Supplementary Fig. 1b). However, when we treated the resulting capsids with Benzonase, we found little to no protection of mRNA (Supplementary Fig.1, c and d).

**Fig. 1.**
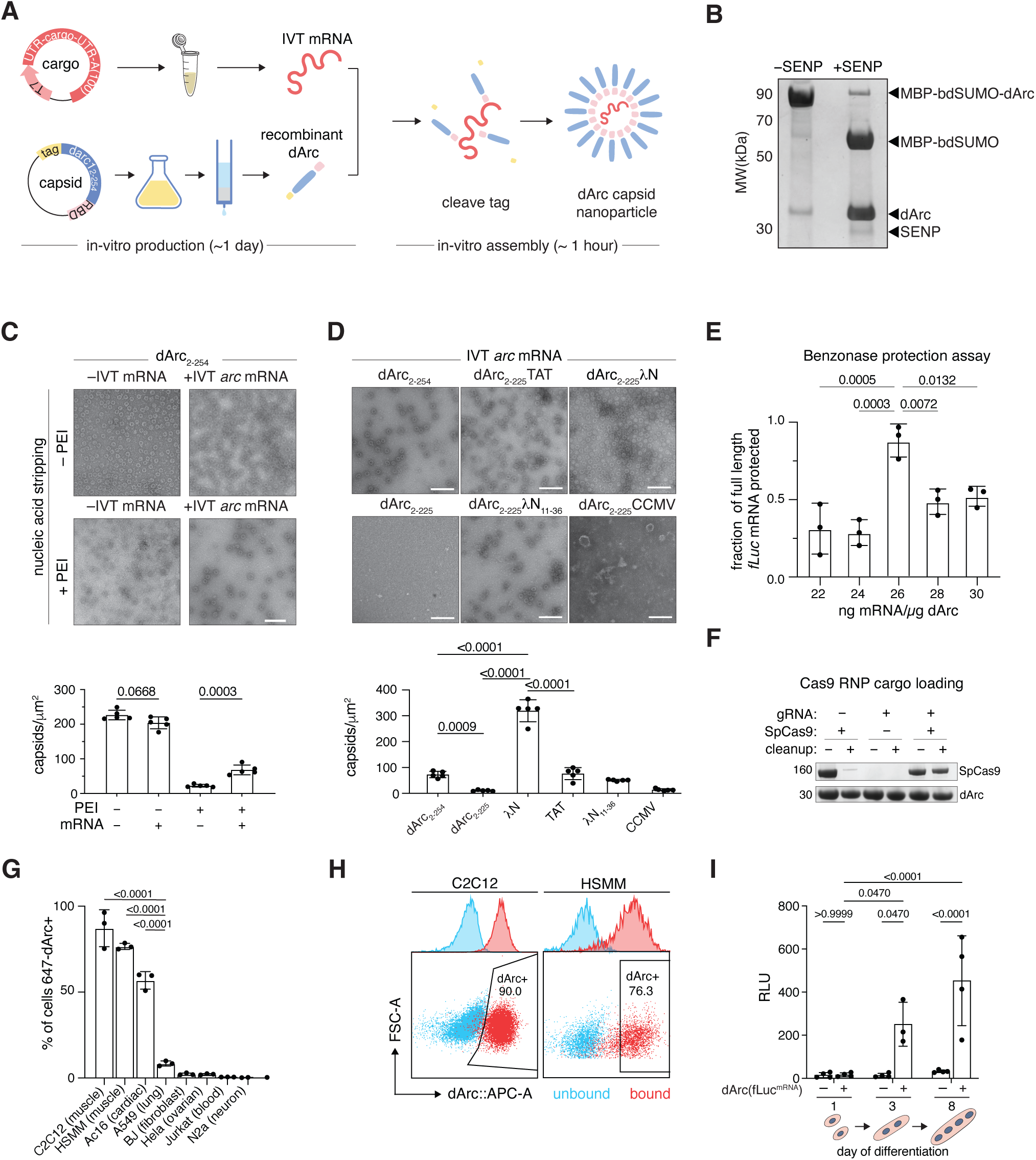
dArc self-assembles into capsids, which can be engineered for cargo delivery *in vitro*. A. Schematic of dArc *in vitro* production and assembly. UTR, untranslated region; IVT, in vitro transcription; RBD, RNA binding domain B. Coomassie staining of a gel showing purified MBP-bdSUMO-dArc before removal of the tag (-SENP) and after removal of the tag (+SENP). MBP, Maltose binding protein; bdSUMO, Brachypodium distachyon SUMO; SENP, Sentrin-specific protease 1 C. (Top) Electron micrographs of dArc capsids either with or without exogenous IVT *arc* mRNA, and with or without nucleic acid stripping with polyethyleneimine (PEI) during protein production at 2.4 mg/mL. (Bottom) Quantification of the number of particles per μm^2^ as determined by electron microscopy. Scale bar represents 200 nm. n=5 fields of view per condition, one-way ANOVA with Tukey’s. D. (Top) Electron micrographs of dArc capsids with modified RNA binding domains, all formulated at 2.4 mg/mL. (Bottom) Quantification of the number of particles per μm^2^ as determined by electron microscopy. Scale bar represents 200 nm. TAT, Trans-Activator of Transcription; CMV, Cowpea Chlorotic Mottle Virus. n=5 fields of view per condition, one-way ANOVA with Tukey’s. E. Quantification of the fraction of input RNA protected by dArc-λN in a Benzonase protection assay with the addition of 640 μM spermidine. One-way ANOVA with Tukey’s, n=3 per condition F. SDS-PAGE gel of dArc capsids pre- and post-Capto Core 400 cleanup G. Quantification of flow cytometry binding of 647-labeled dArc capsids to cell lines of various origins, one-way ANOVA with Tukey’s. n=3 per cell line H. Representative flow cytometry plots of muscle lineage cells binding to 647-labeled dArc capsids. C2C12: murine myoblast cell line; HSMM: Human Skeletal Muscle Myoblasts. I. Functional delivery of *fLuc* mRNA to C2C12 cells at 1, 3, or 8 days of differentiation, two-way ANOVA with Holm-Sidak. n=4

To improve mRNA protection, we focused on engineering the RNA-capsid binding interaction. We generated hybrid capsid fusions wherein the native zinc finger was replaced with RNA-binding domains (RBDs) from various viruses. We found that versions with the bacteriophage lambda N (λN) RBD or the bovine immunodeficiency virus Transactivator of Transcription (BIV-TAT) RBD formed homogenous capsids (Fig. 1d). However, only dArc-λN showed successful nuclease protection, with an optimal RNA:protein ratio of 15-20 ng mRNA per μg of protein (Supplementary Fig.1, d and e). Moreover, the λN variant formed capsids 3-4 fold more efficiently than wild-type dArc (Fig. 1d, Supplementary Fig. 1f). We additionally included an N-terminal SpyTag-002 on the dArc-λN capsid, for downstream external functionalization, which robustly formed capsids (Supplementary Fig. 1g).

Inspired by viral nucleic acid compaction and packaging strategies ^8^, we tested different polyamines during assembly to increase the amount of packaged mRNA per capsid. In a Benzonase protection assay, we found that adding 640 uM spermidine into the packaging reaction increased the total amount of mRNA packaged from 60% to nearly 80% of the input mRNA, and increased the loading ratio of mRNA to protein to an optimum of 26 ng mRNA per μg protein (Fig. 1e, Supplementary Fig. 1h). Using direct RNA nanopore sequencing, we confirmed that after PEI treatment, more than 95% of encapsulated RNA was full-length IVT mRNA (firefly Luciferase, *fLuc*) compared to non-PEI treated capsids (Supplementary Fig. 1i). We measured dArc-λN capsid size by dynamic light scattering (DLS) and found a hydrodynamic radius of ∼39 nm, consistent with the larger apparent size of a fully solvated particle relative to the ∼37 nm diameter observed by EM (Supplementary Fig. 1j). To test if these engineered particles could deliver functional cargo, we treated BJ fibroblasts with dArc-λN capsids packaged with *H2B-mCherry* mRNA (denoted dArc-λN(*H2B-mCherry*^mRNA^)) but did not observe any signal. We hypothesized that this is because the capsids cannot efficiently bind or enter these cells. To test this, we complexed the capsids with cell-penetrating peptides (CPPs) to mediate cell binding and entry (Supplementary Fig. 2a). We screened the CPPs TAT, melittin, LAH4, and R8 and found that the addition of 50 μM LAH4 mediated robust delivery of packaged *H2B-mCherry* mRNA into BJ fibroblasts with minimal background (Supplementary Fig. 2b). Using the LAH4-mediated delivery to BJ cells confirmed that adding 640 uM spermidine into the assembly reaction increased the magnitude of *H2B-mCherry* mRNA delivery by ∼5-fold (Supplementary Fig. 2, c and d) as well as mediated efficient delivery of *fLuc* mRNA at the optimal packaging ratio of 26 ng per μg protein (Supplementary Fig. 2e).

Previous studies have shown that the capsid natively packages its own ∼3-kb mRNA ^1^. In line with this, our engineered dArc-λN capsid can protect 2–3-kb mRNA from nucleases. However, protection and delivery declined substantially after ∼2.5 kb (Supplementary Fig. 3, a and b). Given that the packaging capacity of RNA viruses shows a positive correlation with the charge on the RBD^9^, we reasoned that the charge of the RBD is a limiting factor in packaging larger mRNAs. Thus, we created an additional variant, dArc-SIN, wherein the RBD is swapped for Sindbis virus coat protein 81-113 (Supplementary Fig. 3c), which, like dArc, also forms a T=4 capsid ^10^. This swap increases the net charge on the RBD from +4 (dArc-λN) to +11 (dArc-SIN). We found that dArc-SIN forms homogenous capsids of the same size as dArc-λN and can package and deliver mRNAs of up to at least 5 kb, sufficient to accommodate large genome editors like base and prime editors (Supplementary Fig. 3, c to e).

Next, we tested whether these capsids could package alternative cargo modalities, such as genome editor ribonucleoprotein complexes (RNPs) (Supplementary Fig. 4a), as the transient nature of RNP delivery is less likely to lead to genomic instability and off-target editing. Given that RNA is required for dArc capsid assembly (Fig. 1c), we found minimal packaging and capsid assembly with apo-SpCas9 (Fig. 1f). However, we saw robust capsid assembly not only with sgRNAs alone but also with SpCas9 RNPs. Moreover, we detected SpCas9 protein post-capsid cleanup with Capto Core 400 resin, which removes proteins that are not part of larger complexes, such as capsids (Fig. 1f, Supplementary Fig. 4, b to d). Delivery of dArc-λN(SpCas9^RNP^) with LAH4 led to up to 60% targeted indels in BJ fibroblasts (Supplementary Fig. 4e). These data indicate that engineered dArc capsids can efficiently encapsulate various cargoes and achieve functional cargo delivery.

### dArc capsids bind to cells via interaction with SORCS2

To determine if dArc capsids are capable of autonomous binding and entry into mammalian cells without the need for cationic CPPs, we used capsids packaging a fluorescently-labeled *fLuc* mRNA and screened their binding against a panel of cell lines by flow cytometry. Capsids showed strong binding to C2C12 cells, primary human skeletal myoblasts, and AC16 cardiomyocytes (Fig. 1, g and h, Supplementary Fig.5a), which are all muscle cells. Treatment with proteinase K reduced the fluorescent signal by roughly 60- 70%, indicating that while dArc1 capsids bind to C2C12 cells, only some of the capsids are internalized. (Supplementary Fig.5b). Consistent with this, we found that treatment of C2C12 myoblasts with dArc-λN(*fLuc*^mRNA^) produced minimal luciferase signal in these cells. As myoblasts represent an early state of muscle formation, we next tested if differentiation increases delivery efficiency and observed a pronounced signal, indicating successful delivery of cargo, in differentiating and fully mature myotubes (Fig. 1i).

We next wanted to better understand the molecular mechanism of capsid binding to C2C12 cells. To this end, we generated polyclonal serum against the dArc capsid in mice and performed co-immunoprecipitation mass spectrometry (co-IP/MS) on C2C12 whole cell lysate (Supplementary Fig. 6a). This allowed us to identify two highly enriched membrane proteins, CKAP4 and SORCS2 (Fig. 2a). Given that CKAP4 is primarily localized to the endoplasmic reticulum membrane ^11^, we focused on SORCS2 as the potential binding partner of dArc capsids. We also sorted C2C12 cells treated with capsids packaging fluorescently labeled *fLuc* mRNA based on their fluorescence intensity, indicative of dArc(*fLuc*^647^) binding, into the top 5% and bottom 5% of cells and performed differential gene expression analysis using RNA sequencing. Supporting our co-IP/MS findings, we found *Sorcs2* expression to be significantly enriched in the top 5% (Supplementary Fig. 6b).

**Fig. 2.**
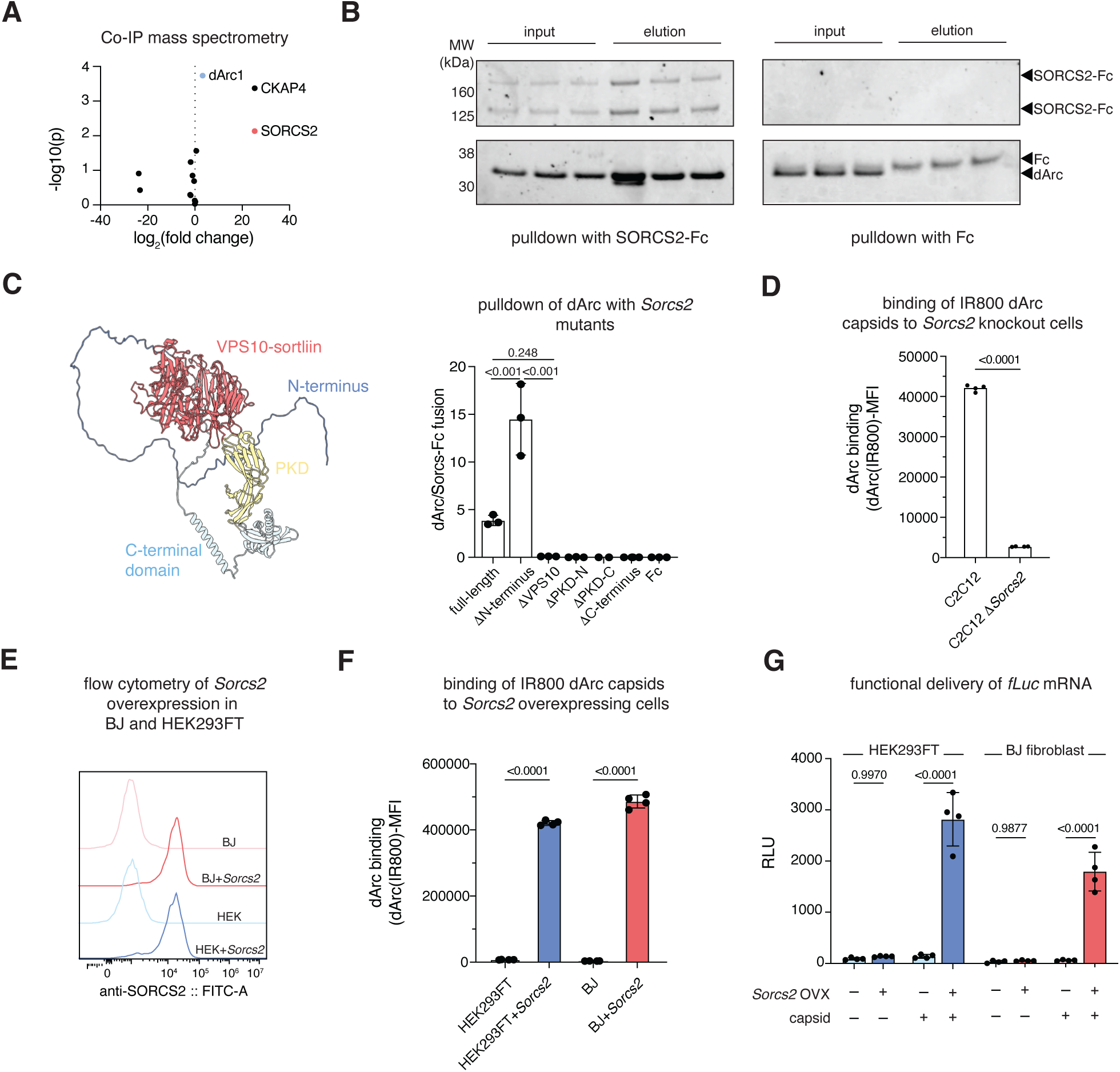
SORCS2 is necessary and sufficient for capsid binding and entry. A. Volcano plot of cell-surface proteins derived from co-IP mass spectrometry of whole cell lysate of C2C12 cells co-precipitated with dArc capsids compared to control IgG pulldown. B. Coomassie stained SDS-PAGE gel of input and elution from pulldowns of dArc capsids with SORCS2-Fc fusion and Fc only. C. Structure of SORCS2 (AlphaFold-DB) and quantification of pulldown of dArc capsids by various SORCS2 N-terminal truncations. N=3 per condition, one-way ANOVA with Tukey’s. D. Mean fluorescence intensity of dArc capsids bound to C2C12 cells or their *Sorcs2* knockout counterpart (C2C12 Δ*Sorcs2*). One-way ANO VA with Sidak. n=4 E. Representative flow cytometry plots of *Sorcs2* overexpressing cell lines compared to parental cell lines stained with an anti-SORCS2 antibody. F. HEK293FT and BJ cells, as well as their Sorcs2 overexpressing counterparts (HEK293FT+Sorcs2 and BJ+Sorcs2) treated with 200 ng IR800-dArc as measured by flow cytometry, one-way ANOVA with Sidak. n=4 G. Functional delivery of *fLuc* mRNA in HEK293FT or BJ fibroblast cells with or without *Sorcs2* overexpression (OVX), two-way ANOVA with Sidak. n=4

To investigate whether SORCS2 directly binds to dArc capsids, we performed a pull-down assay using the ectodomain of murine SORCS2 fused to an Fc fragment (SORCS2-Fc). When incubated with purified dArc capsids, SORCS2-Fc robustly pulled down the capsids. By contrast, the Fc-only control failed to co-precipitate dArc capsids under identical conditions (Fig. 2b). These results demonstrate a direct and specific interaction between the SORCS2 extracellular domain and dArc. SORCS2 is a complex multi-domain type-I transmembrane protein. The ectodomain comprises an N-terminally disordered region, a highly structured VPS-10 sortillin domain (forming a β-propeller), two Polycystic Kidney Disease (PKD) domains, and a structured C-terminal domain with unknown function ^12^. To determine which of these domains is responsible for dArc binding, we immobilized N-terminal truncations of the SORCS2 ectodomain-Fc fusions on beads, and subjected dArc capsids to pulldowns as before. We found that deletion of the β-propeller domain rendered the protein incapable of pulling down dArc capsids (Fig. 2c), indicating that this is the likely interacting domain of dArc capsids on SORCS2. Despite the robust binding of dArc capsids to mammalian SORCS2, high sensitivity structure searches against the *Drosophila* proteome using the VPS10-domain did not reveal any direct orthologs of this domain (Supplementary. 6c).

To functionally interrogate the role of SORCS2 in dArc binding, we generated *Sorcs2* knockout C2C12 cells and observed significantly reduced dArc capsid binding (Fig. 2d), confirming our biochemical results that SORCS2 is essential for the binding of dArc capsids. Conversely, overexpression of *Sorcs2* in cell types where we previously saw no detectable binding, like HEK293FT and BJ fibroblasts (Fig. 2e), led to robust binding (Fig. 2f) and functional delivery (Fig. 2g). These findings show SORCS2 is a critical determinant of dArc capsid delivery and suggest that SORCS2 is both necessary and sufficient to confer permissivity to otherwise non-target cell types.

### Recombinant dArc capsids deliver gene editing payloads *in vivo*

Although *in vitro* we found strong binding and entry to muscle cells due to SORCS2 expression, *in vivo*, *Sorcs2* is expressed highly in the brain, and to a lower extent in many other tissues ^13^ . To characterize the biodistribution of dArc capsids, we first evaluated systemic administration. Intravenously administered capsids carrying fluorescently tagged cargo distributed strongly in the kidney, as well as the liver, spleen, and muscle, but not the brain, consistent with exclusion by the blood-brain barrier (Supplementary Fig. 7a). However, functional delivery was limited, with only low levels of *fLuc* expression detected in the liver (Supplementary Fig. 7, b and c). This is consistent with serum-mediated destabilization of dArc capsids, which showed progressive loss of encapsulated RNA upon serum exposure in vitro (Supplementary Fig. 7, d-f).

Given the observed inhibition in serum, we tested a local delivery approach, by intramuscularly injecting dArc-λN(*fLuc*^mRNA^) capsids into the tibialis anterior (TA) of Balb/c mice and performing bioluminescence imaging 6 hours post-injection. We observed strong bioluminescence signal from the injected muscle, suggesting functional delivery of *fLuc* mRNA into muscle *in vivo* (Fig. 3a, Supplementary Fig.8a and b). As *Sorcs2* expression is modestly upregulated during muscle injury (Supplementary Fig.8c, ^14^), we hypothesized that dArc capsids might show enhanced delivery efficiency in a cardiotoxin-injury model. We induced injury in the TA of wild-type Balb/c mice with cardiotoxin and injected dArc-λN(*fLuc*^mRNA^) in the regenerative phase of injury, when cells damaged from injury have been cleared ^14^ (Day 4) (Supplementary Fig.8d). Bioluminescence imaging 6 hours post-injection showed a ∼10-fold increase in signal intensity for dArc-λN(*fLuc*^mRNA^) capsids compared to uninjured controls (Fig. 3b, Supplementary Fig.8e).

**Fig. 3.**
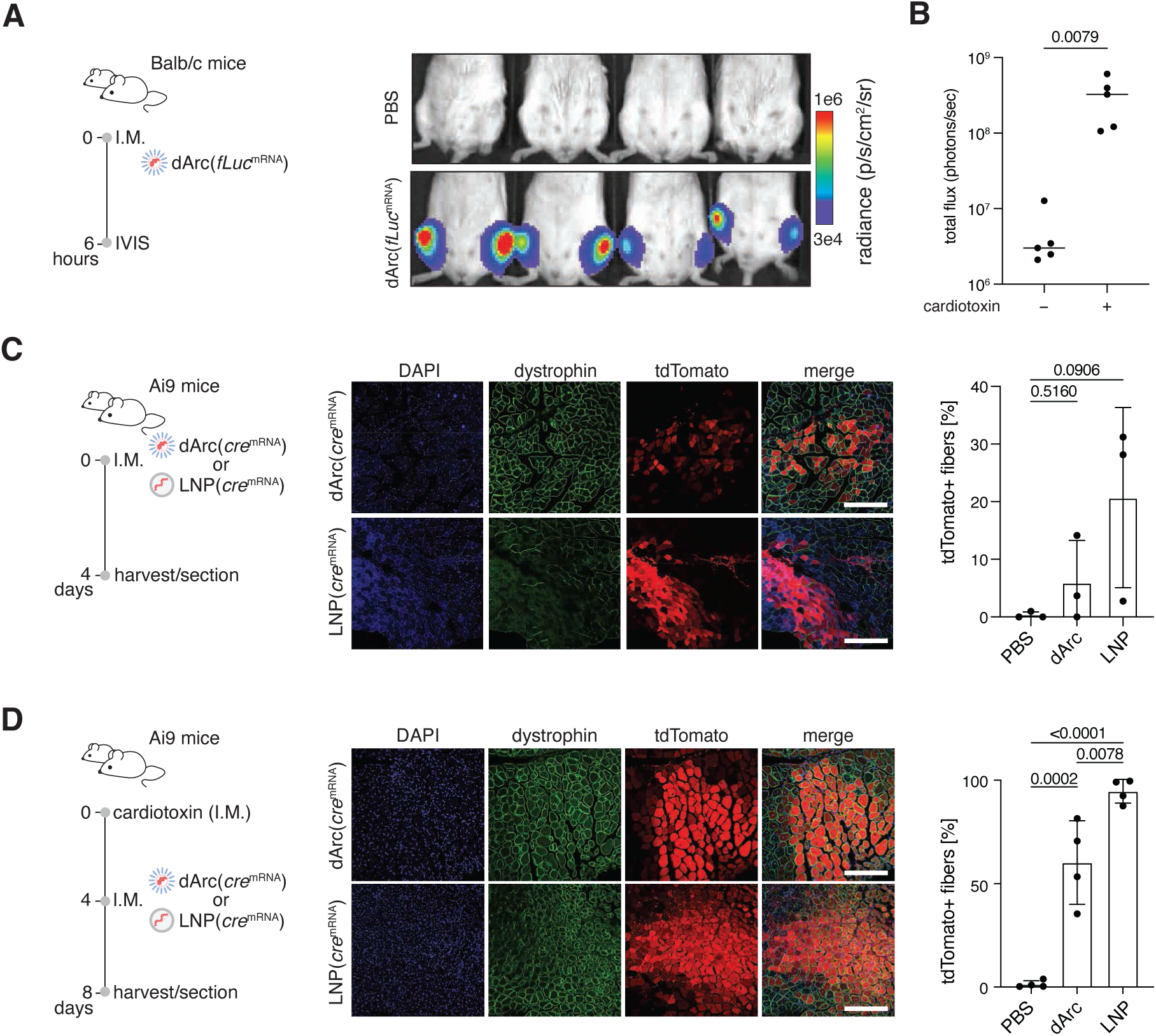
dArc capsids deliver mRNA to muscle cells *in vitro* and *in vivo*. A. Experimental outline (left) and representative bioluminescence imaging (right) 6 hours post-injection of Balb/c mice injected intramuscularly (I.M.) with 4 μg dArc-λN(*fLuc*^mRNA^). B. Quantification of bioluminescence from the cardiotoxin-injured tibialis anterior (TA) and liver of mice injected with 10 μg of dArc-λN(*fLuc*^mRNA^), two-tailed Mann Whitney. C. Experimental design (left), representative immunofluorescence images (middle) and quantification (right) of the TA of mice injected with 10 μg of dArc-λN(*cre*^mRNA^) or 10 μg muscle-tropic TCL053-LNP(*cre*^mRNA^), one-way ANOVA with Tukey’s. The scale bar represents 200 μm. D. Experimental schematic (left), representative immunofluorescence images (middle) and quantification (right) of the TA of mice injected with 10 μg of dArc-λN(*cre*^mRNA^) or 10 μg muscle-tropic TCL053-LNP(*cre*^mRNA^) four days post cardiotoxin injury, one-way ANOVA with Tukey’s. The scale bar represents 200 μm

To confirm that dArc-λN capsids are targeting bona fide muscle cells, we injected dArc-λN(*cre*^mRNA^) particles or positive control muscle-optimized LNPs (TCL053 LNPs) ^15^ encapsulating the matched doses of the same mRNA intramuscularly into Cre reporter mice (Ai9). Sectioning of the injected muscles revealed that dArc-λN(*cre*^mRNA^) and TCL053 LNPs resulted in strong tdTomato expression in muscle fibers, indicative of successful Cre recombination (Fig. 3c). In the cardiotoxin injury model, sectioning of the injected muscle revealed that dArc-λN(*cre*^mRNA^)-mediated delivery was primarily confined to muscle fibers, unlike the LNP control, which appears to have more heterogeneous expression across many cell types (Fig. 3d). In this injury model, dArc particles enable delivery into about 50% of muscle fibers.

As muscle fibers are multi-nucleated, we performed the same experiment in loxP-nuclear-GFP reporter mice to further quantify the degree of muscle fiber vs. non-fiber delivery Supplementary Fig. 8f). Upon nuclear isolation, staining, and flow cytometry we found that cardiotoxin injury increases delivery efficiency to myonuclei, from an average of 8% to 20% (Supplementary Fig. 8g). The percent of mononuclear cells that are GFP+ trends higher with LNP in both settings as compared to dArc (Supplementary Fig. 8i). With dArc particles, cardiotoxin injury increases both the number of positive fibers, as well as the degree of delivery to each fiber (Fig. 3d and Supplementary Fig. 8g). Delivery to hepatocytes was high in the LNP treatment, despite local treatment, reaching nearly 75-80% of hepatocytes in the liver, whereas dArc particles do not mediate delivery to the liver when injected locally (Supplementary Fig. 8h).

Dystrophic muscle undergoes chronic cycles of degeneration and regeneration that resemble the cardiotoxin injury environment ^16^. We therefore tested whether the increased delivery efficiency of dArc-λN(*fLuc*^mRNA^) capsids in injured muscle also translates to dystrophic muscle and found a ∼4-fold increase in luminescence signal when administered to dystrophic mice (mdx-4CV) as compared to age-matched C57BL/6 control mice (Fig. 4a). We hypothesized that the increased signal could be due to targeting of muscle progenitor cells in injured or regenerating muscle, which are known to express SORCS2 ^17^. To test this hypothesis, we crossed dystrophic mice (mdx-4CV) with Cre reporter animals (Ai9) and injected dArc-λN(*cre*^mRNA^) particles intramuscularly into the TA. Analysis by flow cytometry revealed successful delivery of *cre* mRNA to over 20% of all muscle progenitor cells in the TA, indicating that dArc capsids are capable of delivery not only to mature muscle fibers but also to muscle progenitor cells (Fig. 4b, Supplementary Fig. 9a).

**Fig. 4.**
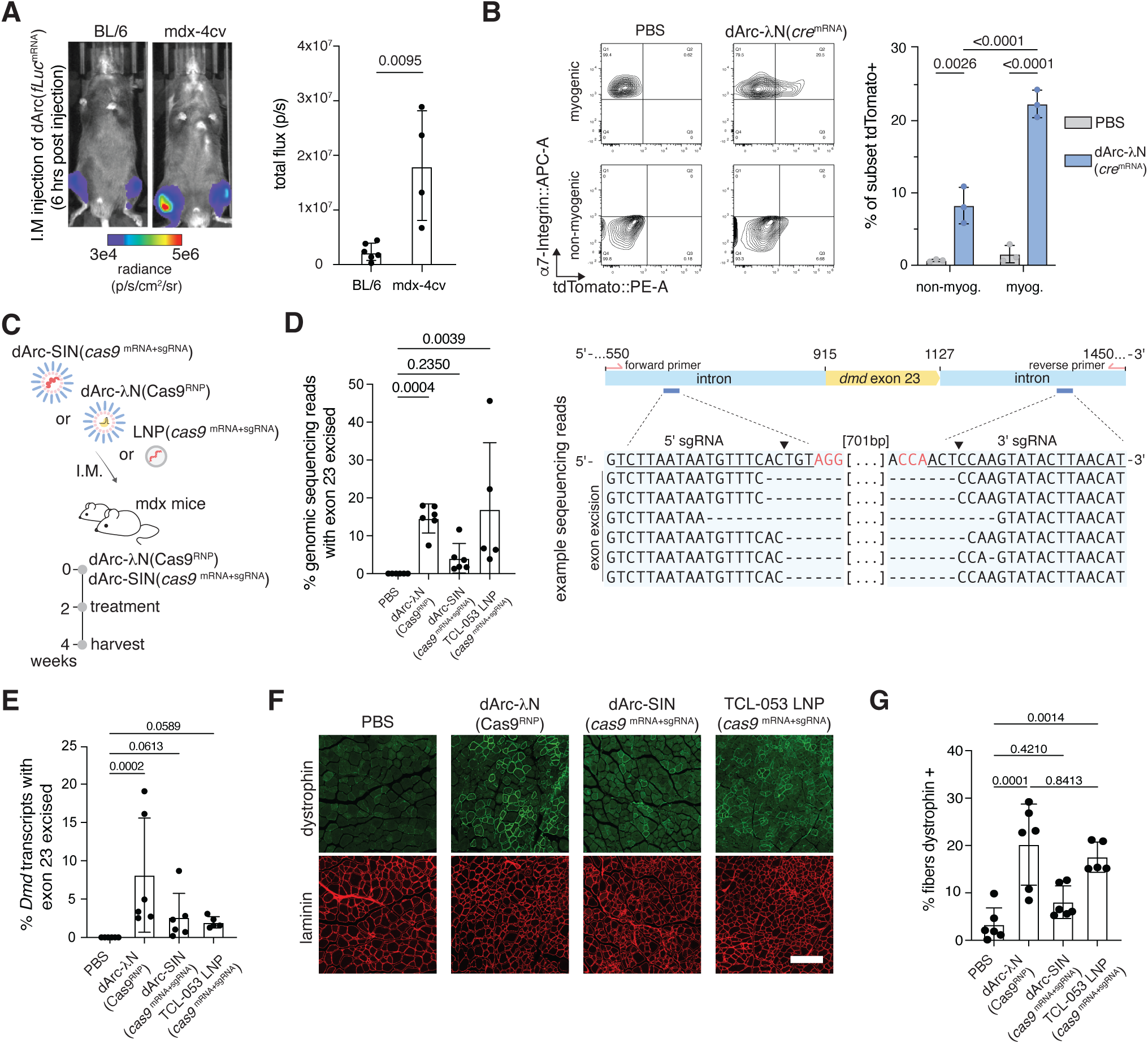
dArc capsids deliver gene-editing cargo to correct *dystrophin* in a mouse model of muscular dystrophy. A. Representative bioluminescence images (left) and quantification (right) 6 hours post-injection of mdx-4CV or age-matched C57BL/6 mice injected I.M. with 4 μg dArc-λN(*fLuc*^mRNA^), n=4 injections, Mann-Whitney test. B. Representative flow cytometry plots (left) and quantification (right) of muscle mononuclear cells dissociated post dArc-λN(*cre*^mRNA^) treatment stained with markers for myogenic and non-myogenic cells. n=3 mice per condition, two-way ANOVA with Tukey’s. C. Schematic of treatment schedule for dystrophin restoration in mdx mice. D. (Left) Quantification of exon excision frequency in genomic DNA and example sequencing reads (right) from mice injected with 10 μg of dArc-λN(Cas9^RNP^) or 10 μg dArc-SIN(*cas9*^mRNA+sgRNA^), Kruskal-Wallis with Dunn’s. E. Quantification of exon excision frequency in mRNA from mice injected with 10 μg of dArc-λN(Cas9^RNP^) or 10 μg dArc-SIN(*cas9*^mRNA+sgRNA^), Kruskal-Wallis with Dunn’s. F. Representative immunofluorescence images of the TA of mdx mice from panels D and E. The scale bar represents 120 μm. G. Quantification of percent of muscle fibers expressing dystrophin from F, one-way ANOVA with Dunnett’s.

To more comprehensively profile on- and off-target cell types, we then performed single-cell RNA sequencing of tdTomato+ mononuclear cells from the muscle of these same mice. Alignment of our data with a previously published single-cell muscle atlas ^18^ indicates delivery to bona fide muscle stem cells and progenitor cells, but also non-myogenic cell types, including monocytes, macrophages, which are part of the mononuclear phagocyte system and are specialized to clear pathogens and debris and thus a common off-target cell type, and fibro-adipogenic progenitor cells, which express *Sorcs2* (Supplementary Fig. 9, b-d). We note that scRNA-seq analysis inherently underrepresents delivery to mature myofibers, as the dissociation protocol required for single-cell suspension destroys multinucleated syncytial fibers, artificially enriching the dataset for mononuclear cell types. The histological analysis (Fig. 3d, 4f) and nuclear flow cytometry data (Supplementary. 8g) provide a more accurate representation of fiber-directed delivery in vivo, demonstrating that most of the dArc-mediated delivery occurs within myofibers. These data demonstrate that dArc capsids retain and enhance their functionality in injured and dystrophic muscle, where SORCS2 upregulation creates a permissive environment for delivery.

Finally, we assessed whether dArc capsids could support the localized delivery of gene-editing tools in a mouse model of muscular dystrophy. We employed a previously described exon excision approach using two optimized paired guides to remove the premature stop codon in exon 23 of the mdx mouse model to restore normal dystrophin expression ^19,20^ (Supplementary Fig. 9, e and f). To test the ability of dArc capsids to deliver multiple types of cargo in the same model system, we formulated dArc-λN(Cas9^RNP^) capsids, or dArc-SIN(*cas9*^mRNA+sgRNA^) capsids, wherein Cas9 mRNA and both sgRNAs were mixed before assembly. We also included a positive control LNP (TCL053 formulation) co-delivering both sgRNAs and *cas9* mRNA, to ensure our optimized sgRNAs worked well *in vivo*. We then injected mdx mice intramuscularly twice with dArc-λN(Cas9^RNP^), dArc-SIN(*cas9*^mRNA+sgRNA^), or LNP (Fig. 4c) and observed exon deletion up to 18% with the RNP strategy (Fig. 4d), with a similar restoration of dystrophin mRNA and protein (Fig. 4, e-g). Cas9 mRNA+sgRNA co-delivery was less efficient, yet still resulted in ∼3.5% exon skipping (Fig. 4, d-g). This is likely due to the added complexities of delivering an mRNA and two sgRNAs into the same cell. Muscle sections of these mice revealed restoration of dystrophin expression in the muscle with both RNP and mRNA+sgRNA delivery (Fig. 4f). Together, these data demonstrate that dArc capsids can deliver therapeutically relevant cargoes and restore dystrophin expression in the muscles of dystrophic mice ^21^ .

## Discussion

In vitro–assembled PNP systems are an attractive option for delivering genetic medicines, given their potential for manufacturing scalability and modular engineering compared to viral vectors ^22,23^. These include systems based on repurposed proteins from lumazine synthase ^24–27^, MS2 bacteriophage ^28–32^, various plant viruses ^33–35^, bacterial proteins ^36,37^, elastin-like polypeptides ^38^, and entirely *in silico-*designed protein nanocages ^39–46^. However, few of these have demonstrated targeted *in vivo* delivery or been adapted for gene editing payloads. dArc is uniquely positioned to address these limitations given its natural capsid-forming activity and amenability to cell-free assembly.

Here we engineer dArc for targeted *in vivo* delivery of multi-modal cargoes, including mRNA and ribonucleoproteins (RNPs). We uncover that dArc capsids bind to mammalian cells, including muscle cells of different species, and that this binding is mediated by SORCS2. While the contribution of additional receptors cannot be excluded, the loss of dArc binding upon SORCS2 knockout in C2C12 cells, combined with the ability of overexpression of SORCS2 to confer functional cargo delivery by dArc in otherwise non-permissive cells, together establish it as a key determinant of cellular uptake. This interaction is particularly surprising, and raises intriguing questions about the identity of the endogenous dArc binding partner in *Drosophila*, which may share an ancient structural fold with SORCS2 that isn’t readily detectable by standard structural search methods. Previous studies identified the linear motif from *Sas* (YDNPSY) as binding to dArc1 ^47^, and there is a similar XPXY sequence in the beta propeller of both human and mouse SORCS2, which may be responsible for binding. Furthermore, dArc functions at the NMJ in Drosophila ^1^ and our data show it can bind and enter mammalian muscle, raising the possibility that its binding partner serves a conserved role at the muscle interface across phyla. We exploited this interaction to deliver cargoes to muscle. We find enhanced delivery in regenerating muscle and differentiated myotubes relative to healthy tissue and myoblasts, respectively, consistent with published transcriptomic data reporting SORCS2 upregulation during cardiotoxin-induced muscle regeneration. Delivery of Cas9 RNP with dArc capsids intramuscularly injected in these animals led to ∼18% dystrophin restoration, within the range of what has been achieved with antisense oligos ^48^.

Despite the advantages of an in vitro assembled PNP for delivery, realizing the full therapeutic potential of this system will require further engineering. First, stabilization of the capsid shell against serum-mediated degradation will be necessary to extend the therapeutic window and enable more efficient systemic administration. Second, endosomal escape remains a key bottleneck for cytosolic cargo delivery. Notably, SORCS2 functions as a sorting receptor that traffics between the plasma membrane and endosomal compartments; understanding whether dArc capsids co-traffic along this route, and at what point cargo is released, may inform rational strategies to enhance escape efficiency across cell types and delivery routes. Finally, it will be important to modify the way in which dArc interacts with human cells to direct its tropism to target tissues. Critically, because dArc capsids can be assembled entirely *in vitro* from recombinant protein and IVT RNA, each component can be independently engineered, characterized, and quality-controlled prior to assembly, affording a degree of compositional precision that is difficult to achieve with viral vectors.

This modularity positions dArc as a uniquely tractable platform for systematic optimization, and establishes a blueprint that may extend beyond dArc itself. The domesticated retroelement capsids found across eukaryotic genomes likely encode a diverse repertoire of tropisms and structural properties, just as the identification of distinct AAV serotypes enabled tissue-specific gene therapies ^49^. Systematic exploration of this natural diversity, combined with the cell-free assembly and engineering principles established here, could substantially expand the toolkit available for therapeutic delivery.

## Data and materials availability

All data are included in the manuscript or supplement.

## Acknowledgments

We acknowledge the Broad Institute Flow Cytometry Core and the MRL EM Core for their technical support. This work was supported in part by the Koch Institute Support (core) Grant P30-CA14051 from the National Cancer Institute. We thank the Koch Institute’s Robert A. Swanson (1969) Biotechnology Center for technical support, specifically Peterson (1957) Nanotechnology Materials Core Facility (RRID:SCR_018674). We also thank James Dahlman, Steve Elledge, and Michael Birnbaum for their valuable insights and all members of the Zhang Lab for feedback and support. B.L. was supported by a National Research Service Award (5F31CA275339-03); F.Z. was supported by the Howard Hughes Medical Institute; the K. Lisa Yang and Hock E. Tan Molecular Therapeutics Center; and the Broad Institute Programmable Therapeutics Donors.

## Author contributions

B.L., D.S., M.S., and F.Z. conceptualized the project. B.L., D.S., M.S., and F.Z. designed the experimental approaches. B.L., D.S., M.S., S.V., K.D., J.P., C.S., P.K., Y.Z., C.L., and J.M. performed experiments. B.L., R.M., and F.Z. acquired funding. R.M. and F.Z supervised the work. B.L. and D.S. wrote the original draft; R.M. and F.Z. revised, reviewed and edited the draft.

## Supplementary Information

$Supplementary Figures 1 – 9

$Tables S1 to S4

Data S1

## Methods

### Cloning and plasmid assembly

Plasmids encoding dArc1 (in Data S1) were all cloned via Gibson assembly. The dArc coding sequence was cloned from a cDNA library generated from total RNA of Drosophila S2 cells. Fragments were generated by PCR using Phusion Flash High-Fidelity 2X Master Mix (Thermo Fisher Scientific Cat# F548L). Fragments were cloned into a 6X-His-MBP-bdSUMO-XTEN backbone using 2X Gibson Assembly master mix (New England Biolabs, Cat# E2611L). The resulting plasmids were transformed into Stbl3 (Thermo Fisher Scientific, Cat# C737303), and correct colonies were identified by next-generation sequencing. Relevant plasmid maps can be found in Data S1, annotated with Plannotate for consistency ^50^.

All mutant plasmids, including additional RNA binding domains, were generated via inverse PCR and ligation using KLD Master Mix (New England Biolabs, Cat# M0554S).

### Protein purification

Protein production plasmids were transformed into Rosetta™ 2(DE3)pLysS (Millipore Sigma, Cat# 71401-4) and grown overnight at 37**°**C. Single colonies were picked for precultures and grown again overnight at 37**°**C in Terrific Broth (TB, USBio, Cat# T2810). On the induction day, cultures were diluted in TB and grown for 6-8 hours at 37**°**C and 220 rpm. Cultures were induced with 0.5 mM IPTG (Gold Bio, Cat# I2481C100) at 18**°**C for 18-20 hours. Cultures were harvested via centrifugation at 4000 xg for 20 minutes at 4**°**C. A standard dArc preparation in our experiments was typically 12 liters of TB, which yields approximately 200-300 mg of nucleic acid and endotoxin-stripped protein (at least 15-25 mg/L yield)

For dArc cultures, pellets were resuspended in lysis buffer (50 mM Tris-HCl pH 8, 400 mM NaCl, 5% glycerol, 2 mM DTT) supplemented with cOmplete ULTRA Tablets (Sigma Aldrich, Cat# 6538282001). The resulting suspensions were lysed on an LM-20 Microfluidizer (Microfluidics) for 2 passes at 18 kpsi. Lysate was clarified via centrifugation at 18,000 x g for 40 minutes and decanted.

For non-nucleic acid-stripped cultures, the clarified lysate was incubated with Ni-NTA Agarose (Qiagen, Cat# 30230) for 30 minutes at 4**°**C. Resin was washed in an Econo Column (Bio-Rad, Cat# 7372532) gravity flow column, washed with 10 CV lysis buffer (without DTT), followed by 10 CV of 1X PBS, pH 7.4 (Gibco, Cat# 10010023). Protein was eluted in PBS with 400 mM imidazole and polished via size exclusion chromatography on a Superdex 200 pg 26/600 column (Cytiva #28989336) in 1X PBS, pH 7.4. Fractions containing the desired protein at >2.4 mg/ml concentration were pooled and frozen at -80 for later use.

For dArc preparations (including dArc-LN, dArc-SIN, etc.) purified with nucleic acid stripping, PEI (Millipore Sigma, Cat# P3143, Mw 750,000) was added to 0.84% into the clarified lysate and mixed via brief inversion. The lysate was immediately centrifuged at 18,000 x g for 30 minutes at 4**°**C. The resulting nucleic acid-stripped supernatant was decanted from the pellet containing other proteins and stripped DNA/RNA. Solid ammonium sulfate (Millipore Sigma, Cat# A4418-5KG) was added to the supernatant while stirring at 4**°**C up to 70% saturation. Protein was left to precipitate for 30 minutes at 4**°**C and then centrifuged at 18,000 x g for 45 minutes. The supernatant was discarded and the protein-containing pellet was resuspended in Butyl-A buffer (50 mM Tris-HCl pH 8, 3M NaCl, 5% glycerol) and filtered through a 0.45 uM Stericup filter (Millipore Sigma, Cat# S2HVU02RE). Protein was applied to a custom-packed 185 mL Butyl Sepharose HP column (Cytiva, Cat# 17-5432-02) and washed with 5 CV Butyl-A buffer. Protein was eluted in a 2 CV gradient elution from 0 to 100% Butyl-B (50 Tris-HCl pH 8, 5% glycerol). Fractions containing the desired protein were collected and applied to a HisPrep FF 16/10 (Cytiva, Cat# 28-9365-51) column. The column was washed with 10 CV IMAC wash (50 mM Tris-HCl pH 8, 400 mM NaCl, 5% glycerol), 15 CV of endotoxin stripping buffer (50 mM Tris-HCl pH 8, 400 mM NaCl, 5% glycerol, 0.1% Triton X-114), 5 CV IMAC Wash, 20 CV PBS with 20 mM imidazole, and eluted via 5CV gradient elution in PBS + 400 mM imidazole. Protein-containing fractions were pooled and polished on a Superdex 200 pg 26/600 column in 1X PBS.

In a typical 12-liter dArc preparation, the elutions from the Butyl column were split into two and applied in two successive runs to HisPrep FF 16/10 columns to prevent protein loss. From these two IMAC runs, we typically collect between 45-60 mL of elution, which is loaded in 3-4 successive Superdex 200 pg 26/600 runs at 15 mL per run. After size exclusion, fractions with a concentration greater than or equal to 2.4 mg/mL are pooled and frozen at -80**°**C for later use. We have found that MBP-bdSUMO-XTEN-dArc protein irreversibly precipitates if concentrated via centrifugal filters. This protocol avoids this, and we strongly recommend that this protein not be concentrated via centrifugation. Representative chromatograms can be found in Supplementary Fig.1A.

### In-vitro transcription of mRNA

For all mRNAs, except the dArc mRNAs used in Fig. 1, coding sequences were cloned into a custom-built vector containing a T7 promoter, human alpha-globin 5’UTR, a synthetic 3’ UTR, and a A70LA30 interrupted polyA tail. Vectors were designed with a PsiIv2 restriction enzyme site at the 3’ end. Plasmids were cloned and midi-prepped with the ZymoPURE II Plasmid Midiprep Kit (Zymo Research #D4201). Templates were digested with PsiIv2 (New England Biolabs #R0744) per manufacturer instructions. Linearized templates were purified with Ampure XP and used as input for an IVT reaction. For dArc mRNAs, templates were created using PCR of a CMV overexpression vector containing the dArc1 mRNA sequence (CDS and 3’ UTR). IVT mRNA was generated using the NEB HiScribe™ T7 High Yield RNA Synthesis Kit (NEB #E2040L), following modified manufacturer instructions. Reaction buffer was used at 0.5X, nucleotides at 5 mM final concentration, and 2 µg of linearized template was used per reaction. Capping was done co-transcriptionally by the addition of CleanCap® Reagent AG (Trilink Biotechnologies #N-7113) into the reaction, and uridine was completely substituted with N1-Methylpseudouridine-5’-Triphosphate (Trilink Biotechnologies #N-1081). Reactions were conducted at 37**°**C for 5 hours and purified using lithium chloride (Thermo Fisher #AM9480) precipitation. mRNA for *in vivo* use was additionally purified via cellulose purification, as previously described ^51^. mRNA was resuspended, aliquoted, and frozen at -80**°**C in 1 mM sodium citrate, pH 6.5 (Thermo Fisher #AM7001).

For the generation of fluorescent mRNAs, IVT reactions were performed as described, except uridine was substituted for 80% N1-Methylpseudouridine-5’-Triphosphate and 20% 5-DBCO-PEG4-UTP (Jena Bioscience #CLK-055). mRNAs were purified with lithium chloride precipitation, resuspended in water, and reacted with 5-fold molar excess of azide-fluorophore overnight at 4**°**C, either AZDye 647 Azide (Vector Labs #CCT-1299) or AZDye 800 Azide (Vector Labs #CCT-1562) for *in vitro* and *in vivo* use. mRNAs were purified again with lithium chloride precipitation and extensively washed in ice-cold 70% ethanol to remove excess dye. mRNA was resuspended, aliquoted, and frozen at -80**°**C in 1mM sodium citrate, pH 6.5 (Thermo Fisher #AM7001). All mRNAs were quantified with Nanodrop.

### Capsid assembly reactions

Capsid assembly reactions were performed in 1X PBS, pH 7.4 at a 2.4 mg/mL protein concentration. Aliquoted protein and IVT mRNA were thawed on ice. A 64 mM stock of spermidine (Millipore Sigma, Cat# 85558) was made in water. mRNA was first denatured at 65**°**C for 5 minutes before assembly. To assemble capsids, an amount of mRNA corresponding to the desired amount of protein (using an optimized ratio of 26 ng mRNA to ug of protein) was transferred into a tube. A corresponding amount of spermidine is added on ice, immediately followed by MBP-bdSUMO-dArc protein in PBS. This was mixed well and allowed to assemble on ice for 10 minutes. After this wait time, bdSENP1 (produced in house) was added to a 10 µg/mL final concentration. This assembly and cleavage reaction was allowed to sit on ice for at least 1 hour until further processing.

For *in vitro* use, capsids were purified via multi-modal chromatography over CaptoCore 400 resin (Cytiva, Cat# 17372402). Briefly, the 50% stock slurry was equilibrated in PBS via centrifugation and resuspension. For each 100 ul of capsid assembly reaction, 200 ul of 50% slurry was added to a 96-well chromatography plate (Harvard Apparatus, Cat# 4-5650). PBS was removed by centrifugation, and 100 ul of capsid assembly was added per well. The plate was shaken at 400 rpm for 10 minutes at room temperature and then centrifuged at 1200 x g for 5 minutes. The flow-through contains the clean and assembled capsids.

For *in vivo* use, capsids were purified via ultracentrifugation. Capsid assembly reactions were spun in a Beckman Coulter Optima XPN-80 ultracentrifuge equipped with an SW28 rotor for 2 hours at 26,000 rpm over a 20% sucrose cushion. The supernatant was decanted, the pellet dried briefly and resuspended in 1X PBS for use *in vivo*.

Loaded RNA was quantified using the Quant-it™ RiboGreen RNA Assay Kit (Thermo Fisher Scientific, Cat# R11490) per manufacturer instructions. We found the capsids to be permeable to RiboGreen reagent, and thus no previous processing is necessary to quantify mRNA or sgRNA in this manner. To quantify loaded protein, 2 ul of unconcentrated capsid product was loaded on an SDS-PAGE gel alongside a standard curve of the protein of interest (e.g. SpCas9). Gels were stained with Coomassie Blue and imaged on a BioRad ChemiDoc. Encapsulation of proteins was determined by comparing the amount of protein pre and post-cleanup to a standard curve using BioRad ImageLab (https://www.bio-rad.com/en-us/product/image-lab-software?ID=KRE6P5E8Z). All dosing information in this paper is with respect to the mRNA or sgRNA, 10 ug of dArc(mRNA) refers to 10 ug mRNA, whereas 10 ug dArc (RNP) refers to 10 ug sgRNA and the matching amount of Cas9.

### Electron microscopy

Carbon-coated 200-mesh copper grids (Electron Microscopy Sciences, Cat# CF200-CU-50) were glow discharged for 1 minute. Samples in PBS were applied in 5 ul for 1 minute, followed by washing in 5 drops of 2% uranyl formate (Electron Microscopy Sciences, Cat# 22451) for 5, 5, 10, 20 and 20 seconds, respectively. Grids were blotted dry and imaged on an FEI Tecnai transmission electron microscope at 120 kV.

To quantify capsid density from electron micrographs, a particle picking class averaging approach was used to separate capsids from non-capsids. Particles were picked using the standard negative stain model and parameters in crYOLO ^52^. These particles and micrographs were imported into RELION 5.0 (https://github.com/3dem/relion), and class averages were calculated. Particles were classified into capsid and non-capsid classes based on the class averages.

### Benzonase protection assays

Benzonase protection assays were performed immediately after capsid assembly but before purification. Post-assembly, magnesium chloride (Thermo Fisher Scientific, Cat# AM9530G) was added to the reaction to 1 mM, followed by 1U/ul of benzonase (Sigma Aldrich, Cat# E1014-25KU). Digestion was allowed to take place for 10 minutes at room temperature, followed by purification using Capto Core 400 as described in “Capsid assembly reactions”. 5 ul of the pre-benzonase and 5 ul of post-benzonase capsids were extracted using Trizol and the Direct-zol 96 RNA Extraction Kit (Zymo Research, Cat# R2057). RNA was eluted in 25 ul per well and 5 ul of the eluted RNA was run on a 1% 48-well E-Gel (Thermo Fisher Scientific, Cat# G800801) and imaged on a BioRad ChemiDoc.

### Cell culture

BJ fibroblasts, C2C12, A549, HeLa, and N2a were obtained from ATCC and cultured in DMEM (Gibco, Cat# 11965092) + 10% FBS (VWR Seradigm, Cat# 97068-085) + 1% penicillin-streptomycin (Gibco, Cat# 15-140-122). HEK293FT (Thermo Fisher Scientific, Cat# R70007) cells were cultured in DMEM + 10% FBS + 1% penicillin-streptomycin. AC16 cells were obtained from Sigma-Aldrich (Cat# SCC109) and cultured in DMEM:F12 (Gibco, Cat# 10565018) + 12.5% FBS + 1% penicillin-streptomycin. Jurkat E6 cells were obtained from ATCC (Cat# TIB152) and cultured in RPMI 1640 (Gibco, Cat# 61870036) + 10% FBS + 1% penicillin-streptomycin. Human skeletal muscle myoblasts (HSMM) were obtained from Lonza (Cat# CC-2580) and cultured with the SkGM-2 Skeletal Muscle Cell Growth Medium-2 BulletKit (Lonza, Cat# CC-3245).

### Functional delivery with LAH4

For functional delivery with LAH4, LAH4 peptide was purchased from Genscript (RP20096) and resuspended in DMSO (Millipore Sigma, Cat# D2650) at 20 mM. For delivery experiments, LAH4 was added at the appropriate concentration to the capsid mixture, allowed to incubate for 5 minutes at room temperature, and the mix was then added to cells as needed.

### RNP capsid assembly reactions

SpCas9 RNPs were assembled as previously described. Chemically modified Alt-R sgRNAs were synthesized by IDT and resuspended at 1000 ng/ul in water. Equimolar amount of SpCas9 (IDT, Cat# 1081059) and sgRNA were mixed and allowed to form RNPs at room temperature for 10 minutes. An amount of MBP-bdSUMO-dArc protein is added to the preformed RNP at a ratio of 26 ng of sgRNA per µg of MBP-bdSUMO-dArc protein. This assembly reaction is allowed to take place for 10 minutes on ice, and is then processed and cleaned as described above in “Capsid assembly reactions”.

### NGS for indel sequencing (in-vitro)

Genomic DNA from cells in gene editing experiments was extracted with QuickExtract DNA Extraction Solution (Biosearch Technologies, Cat# QE09050). Extract corresponding to at least 10,000 cells was used as input to a PCR using target specific oligos (Table 1) using Phusion Flash High Fidelity 2X Master Mix (Thermo Fisher Scientific, Cat# F548L). This extract was amplified for 15 cycles, and then 1 ul of this PCR was used as input to a barcoding PCR to ligate Illumina adapters and barcodes for sequencing. Libraries were pooled and purified using Ampure XP (Beckman Coulter, Cat# A63881) and sequenced on an Illumina MiSeq. Indels were quantified using a Matlab script ^53^.

### Cell binding assays

For binding assays, we generated fluorescent capsids by encapsulating 647 or IR800-labeled fLuc mRNA. Adherent cells were first washed with PBS and lifted with TrypLE (Thermo Fisher Scientific, Cat# 12604-021) or ReLeSr (Stemcell Technologies #100-0484) to generate single cell suspensions. 200 ng of capsid was added to 100,000 cells in 50 ul of FACS buffer (PBS+3% FBS+2mM EDTA) per well. Binding was allowed to occur at 4C for 1 hour, after which cells were washed 3 times in 200 ul of FACS buffer. If indicated, cells were proteinase K (50 μg/ml) for 30 minutes at 37C and washed again. Cells were resuspended in 100 ul FACS buffer + DAPI for flow cytometry. Flow cytometry for cell binding assays was performed on a CytoFlex-S (Beckman Coulter) or a Sony ID7000 Spectral Cell Analyzer.

### Cell line generation

Mouse *Sorcs2* was cloned from C2C12 cDNA into pLVX-IRES-Puro, pLVX-EF1alpha-SARS-CoV-2-N-2xStrep-IRES-Puro was a gift from Nevan Krogan (Addgene plasmid # 141391. psPAX2 was a gift from Didier Trono (Addgene plasmid # 12260). pMD2.G was a gift from Didier Trono (Addgene plasmid # 12259). Lentiviruses were produced as previously described^54^. HEK293FT cells were transfected with envelope, lentiviral helper plasmid, and one of the above transfer genomes using PEI (Polysciences, Cat# 24765). Four hours later, media was exchanged for fresh DMEM+10% FBS+1% PenStrep. 2 days post transfection, media was harvested, clarified via centrifugation at 1500xg for 5 minutes, and filtered through a 0.45-µm filter (Millipore Sigma, Cat. # SE1M003M00). For cell line generation, C2C12 cells were seeded at 500,000 cells per well in a 6 well plate. 2 mL of unconcentrated clarified supernatant was added to each well 24 hours after seeding, and transduction was allowed to occur overnight. Media was exchanged the next day and cells were returned to the incubator for an additional 24 hours. 48 hours after transduction, cells were selected with 0.5-1 ug/mL puromycin (Gibco, Cat# A1113803) for 5-6 days before use.

### Immunoprecipitation experiments

To generate polyclonal sera against dArc capsids, dArc(fLuc) was injected twice into 6-8 week old Balb/c mice with two week spacing. Two weeks after the last dose, mice were euthanized and blood collected via cardiac puncture into serum separator tubes (BD 365967) and serum was collected per manufacturers instructions. Serum was heat inactivated at 55C for 30 minutes, then aliquoted and stored at -20C. Control serum was collected from PBS injected Balb/c mice using the same workflow. To generate resin for use in immunoprecipitation experiments, antibodies were bound and crosslinked to agarose beads using the Pierce Crosslink IP kit (Thermo Fisher Scientific Cat.# 26147). After following manufacturers instructions for antibody crosslinking, 750 ng of dArc(Cre) capsid were bound to the beads overnight with end over end mixing. The next morning, resin was washed per kit instructions. Whole cell lysate was generated by lysing C2C12 cells in RIPA buffer (Thermo Fisher Scientific Cat.# 89900) + 0.5 % Triton-X100 for 30 minutes on ice, followed by a clarification spin at 15,000xg for 30 minutes at 4C. Protein was quantified using the Qubit™ Protein assay kit (Thermo Fisher Scientific Cat.# Q33211), and 500 ug of whole cell lysate was added to each column and allowed to bind overnight. The following morning the columns were washed and eluted according to the Pierce Crosslink IP kit instructions, and eluates were neutralized by addition of Tris-HCl, pH 8.0 to 50 mM.

### Mass spectrometry

#### Digestion for S-trap samples

Samples were digested on S-trap micro spin columns from Protifi. Samples were digested following the manufacturer’s protocol with slight changes.

Step 5: 10 mM DTT (final concentration) was used instead of TCEP to reduce the proteins and the samples were placed on a heating block for 10 minutes at 95C.

Step 6: 20mM iodoacetamide (final concentration) was used instead of MMTS, and samples incubate at RT for 30 minutes in the dark.

#### LC-MS

The tryptic peptides were separated by reverse phase HPLC (Thermo Ultimate 3000) using a Thermo PepMap RSLC C18 column (2um tip, 75umx50cm PN# ES903) over a 100 minute gradient before nano electrospray using a Orbitrap Exploris 480 mass spectrometer (Thermo). Solvent A was 0.1% formic acid in water and solvent B was 0.1% formic acid in acetonitrile.Solvent A was 0.1% formic acid in water and solvent B was 0.1% formic acid in acetonitrile.

#### Gradient

| Time | % B buffer | Flow rate nL |
| --- | --- | --- |
| 0 | 1 | 300 |
| 20 | 1 | 300 |
| 25 | 1 | 200 |
| 30 | 7 | 200 |
| 65 | 25 | 200 |
| 75 | 36 | 200 |
| 75.5 | 80 | 200 |
| 80 | 80 | 200 |
| 80.1 | 1 | 200 |
| 100 | 1 | 200 |

The Thermo Orbitrap Exploris 480 mass spectrometer was operated in a data-dependent mode. The parameters for the full scan MS were: resolution of 120,000 across 375-1600 *m/z* and maximum IT 25 ms. The full MS scan was followed by MS/MS for as many precursor ions in a two second cycle with a NCE of 28, dynamic exclusion of 20 s and resolution of 30,000.

#### Database search with Proteome Discoverer

Raw mass spectral data files (.raw) were searched using Sequest HT in Proteome Discoverer (Thermo). Sequest search parameters were: 10 ppm mass tolerance for precursor ions; 0.05 Da for fragment ion mass tolerance; 2 missed cleavages of trypsin; fixed modification was carbamidomethylation of cysteine; variable modifications were oxidation of Methionine, Methionine loss at the N-terminus of the protein, acetylation of the N-terminus of the protein, Met-loss plus acetylation of the protein N-terminus. Data was searched against a Uniprot Mouse database alongside sequences for the dArc1 capsid, MBP-bdSUMO tag, and a contaminate database made in house.

#### SORCS2 pulldown experiments

Mouse *Sorcs2* was subcloned into a pCMV-Twist vector fused to a human IgG1 Fc domain via Gibson assembly. Mutants were cloned from this base plasmid using inverse PCR followed by ligation. All SORCS2 fusion proteins were produced by transient transfection of HEK293FT cells using PEI (Polysciences, Cat# 24765). Two days post transfection, supernatant was clarified via centrifugation at 1500xg for 5 minutes, and filtered through a 0.45-µm filter (Millipore Sigma, Cat. # SE1M003M00). Supernatant was mixed with Pierce Protein A Magnetic Beads (Thermo Fisher Scientific Cat.# 88845) and protein was bound to beads overnight at 4C with mixing. The next day, beads were washed three times with PBS+0.1% Tween 20 and used to pulldown dArc capsids as described in “Immunoprecipitation experiments”.

#### Animal experiments

All experiments were performed in accordance with protocols approved by the Institutional Animal Care and Use Committee (IACUC) of the Broad Institute (Protocol #0017-09-14-3). All animals were 6-10 weeks of age at time of experiment. Mice were obtained from either Charles River Laboratories for Balb/c (Strain Code 028) or Jackson Laboratories for C57BL/6J (Strain #:000664), mdx (Strain #:001801), mdx-4CV (Strain #:002378), Ai9 (Strain #:007909), CAG-Sun1/sfGFP (loxP-nuclear-GFP, Strain #021039). Ai9 x mdx-4CV crossings were generated by mating mdx-4CV females with Ai9 males, hemizygous mdx-4CV/heterozygous Ai9 males generated from this mating were used for the experiments in this paper as dystrophin is X-linked.

#### Systemic injections

##### For all systemic injections

capsids or LNPs were suspended in 100 ul of sterile saline and injected into retrorbitally under isoflurane (Covetrus North America, Cat# 029405) anesthesia. Mice were allowed to recover until ambulatory and monitored until harvest. For fluorescent biodistribution studies, animals were euthanized at 6 hours post injection, perfused with PBS, and organs were weighed and then imaged using a BioRad GelDoc. Quantification of MFI from imaged organs was performed on FIJI/ImageJ (https://imagej.net/software/fiji/downloads).

For bioluminescence experiments, mice were anaesthetized with isoflurane and injected with IVIS-brite D-luciferin (Revvity, Cat# 122799) in PBS at 150 mg/kg subcutaneously. Ten minutes post-injection mice were imaged on an IVIS Spectrum CT (Revvity) using automated exposure settings and euthanized immediately after. Bioluminesence was quantified using Living Image software (Revvity, https://www.revvity.com/software-downloads/in-vivo-imaging).

#### Intramuscular injections

For all intramuscular injections, capsids or LNPs were suspended in 40 ul of sterile saline and injected into the tibialis anterior (TA) under isoflurane (Covetrus North America, Cat# 029405) anesthesia. Mice were allowed to recover until ambulatory and monitored once daily for seven days post injection. For cardiotoxin injections, cardiotoxin (Millipore Sigma #217503-1MG) was injected as previously described ^55^. Mice received a single dose of meloxicam (Patterson Veterinary Supply, Cat# 78937565) subcutaneously at 5 mg/kg before cardiotoxin injection. 35 ul of 10 uM cardiotoxin in sterile saline was injected intramuscularly into the TA under isoflurane anaesthesia. Mice were allowed to recover and monitored daily for mobility deficiencies. Four days post cardiotoxin injury, capsids or LNPs were injected for relevant experiments.

Mice were euthanized through CO_2_ asphyxiation and cervical dislocation. For experiments involving Ai9 mice, mice were transcardially perfused with 4% paraformaldehyde (PFA) in PBS and muscles were harvested and post-fixed in 4% PFA (Electron Microscopy Services, Cat# 15710-S) for 2 hours. Muscles were dehydrated in 30% sucrose solution and embedded in Tissue-Tek® O.C.T. Compound (Sakura, Cat# 4583) by snap freezing in liquid nitrogen. For experiments involving mdx mice, mice were transcardially perfused with PBS and muscles were harvested and dissected. Biopsies were taken and snap-frozen in liquid nitrogen for downstream molecular analysis, and the remainder of the tissue was fixed in 4% paraformaldehyde in PBS overnight at 4C. Muscles were dehydrated in 30% sucrose solution and embedded in Tissue-Tek® O.C.T. Compound by snap freezing in liquid nitrogen.

#### Muscle tissue sectioning and immunofluorescence

For all experiments, muscles were sectioned transversely in 12 μM sections on a Leica CM1950 cryostat at -16C. Sections were mounted on VWR® Premium Superfrost® Plus Microscope Slides (VWR #48311-703) and dried at room temperature for one hour before freezing at -80C. Before staining, slides were brought to room temperature. Slides were washed and permeabilized in PBS + 0.1% Triton X-100 (Sigma-Aldrich, Cat# 93443) for 15 minutes at room temperature. Sections were blocked and stained in PBS + 0.1% Triton X-100 + 5% normal donkey serum (Abcam, Cat# ab7475) overnight at 4C. Sections were stained with rabbit anti-dystrophin (Abcam, Cat# ab15277, 1:300) and rat anti-laminin (Thermo Fisher Scientific Cat.% MA1-06100) where indicated. Sections were washed three times in PBS + 0.1% Triton X-100 before staining with Alexa Fluor 488 donkey anti-rabbit IgG (Thermo Fisher Scientific, Cat# A-21206), and Alexa Fluor 594 donkey anti-rat IgG (Thermo Fisher Scientific, Cat# A-21209). Slides were again washed three times and mounted in VECTASHIELD Vibrance® Antifade Mounting Medium with DAPI (Vector Laboratories, Cat# H-1800). Slides were dried and imaged on a Leica Stellaris 5 microscope.

#### Muscle progenitor analysis

To analyze and analyze or sort muscle mono-nuclear cells, mice were euthanized through CO_2_ asphyxiation and cervical dislocation. The TAs from each mouse were harvested and pooled. Single-cell suspensions were generated using the Skeletal Muscle Dissociation Kit, mouse and rat (Miltenyi Biotec, #130-098-305) using a GentleMACS Octo-Dissociator with Heaters (Miltenyi Biotec). Single-cell suspensions were filtered through 70 uM filters and spun down and resuspended in FACS buffer (PBS + 3% FBS + 2mM EDTA). Cells were Fc blocked using TruStain FcX™ (anti-mouse CD16/32) (Biolegend, Cat#101320) for 20 minutes at 4C. Cells were then stained for 30 minutes in FACS buffer with a lineage cocktail, FITC anti-mouse Ly-6A/E (Biolegend #108105), FITC anti-mouse TER-119 (Biolegend #116205), FITC anti-mouse CD31 (Biolegend #102405), and FITC anti-mouse CD45 (Biolegend #103107) as well as ITGA7 Monoclonal Antibody (334908), APC (Thermo Fisher #MA5-23555) all at 1:200. Cells were washed three times in FACS buffer and resuspended in FACS buffer with DAPI for analysis and sorting. Cells were analyzed and sorted on a Sony MA-900 cell sorter.

#### Nuclear isolation and flow cytometry

To isolate nuclei from muscle and liver, tissue was minced into small pieces using scissors, and homogenized in buffer (250 mM sucrose, 10 mM Tris-HCl (pH 8.0), 25 mM KCl, 5 mM MgCl_2_ by 20 strokes in a dounce homogenizer. Triton-X100 was added to this mixture to a final concentration of 0.1%, mixed, and allowed to lyse for 5 minutes on ice. Crude homogenate was filtered through a 70 uM strainer and spun down at 3000xg for 10 minutes. Pellets were resuspended in FACS buffer (PBS + 3% FBS + 2mM EDTA) and filtered again through a 30 uM cell strainer. The resulting nuclei were stained in FACS buffer using an anti-PCM-1 antibody ^56^ (for myonuclei; Millipore Sigma Cat.# HPA023374-100UL) at a 1:100 dilution in FACS buffer, or an anti-HNF4A antibody (for hepatocyte nuclei; Thermo Fisher Scientific Cat.# MA5-14891) at 1:100 in FACS buffer for 1 hour at 4C. Nuclei were washed three times in FACS buffer and stained with Alexa Fluor 488 donkey anti-rabbit IgG (Thermo Fisher Scientific, Cat# A-21206) at 1:300 in FACS buffer for 30 minutes. Nuclei were again washed three times and resuspended with FACS buffer + DAPI. Nuclei were analyzed on a Beckman Coulter Cytoflex-S flow cytometer.

#### Single-cell sequencing

Cells were sorted into FACS buffer, spun down at 1000xg, and resuspended in PBS for input into single-cell library preparation. Single-cell libraries were prepared using Chromium GEM-X Single Cell 3’ Kit v4 (10X Genomics, Cat# 1000686) processed with a Chromium-X controller. Libraries were prepared according to manufacturer instructions and sequenced as recommended on an Illumina NextSeq550 150 cycle high-output kit (Illumina #20024907). Data was demultiplexed and processed with cellranger (10X Genomics, https://www.10xgenomics.com/support/software/cell-ranger/latest) and aligned to the mm10 transcriptome to be compatible with previously published atlas datasets ^22^. Sequenced cells were integrated into this previous atlas dataset using Harmony and Seurat.

#### Muscle molecular analysis (qPCR, NGS)

For DNA extraction, frozen muscle tissue was thawed and DNA was extracted using the DNAdvance Kit (Beckman Coulter, Cat# A48705). DNA was eluted as recommended in the manufacturer’s instructions and 1 ul was used as input to a PCR using gene specific primers with Phusion Flash Master Mix. Samples were cycled 15 times using these primers, and then 1 ul was used as input to a barcoding PCR to add multiplexing barcodes and Illumina adapters for sequencing. Amplicons were cleaned up using AmpureXP and sequenced on an Illumina MiSeq. Reads were demultiplexed and analyzed in Python.

For RNA extraction, frozen muscle tissue was pulverized using a MultiSample BioPulverizer (Biospec Products Cat. No. 59012MS) under liquid nitrogen temperatures. Pulverized tissue was used as input to the RNAdvance Tissue Kit (Beckman Coulter, Cat# A32649) per manufacturer instructions. Eluted RNA was used as input to RT-qPCR using TaqPath™ 1-Step RT-qPCR Master Mix (Thermo Fisher Scientific, Cat# A15299) and previously validated primers and probes against the mouse dystrophin exon 4-5 junction, and the non-canonical exon 22-24 junction created by exon excision ^23^. A primer set against the 18S rRNA was used as a housekeeping control to normalize between samples (Thermo Fisher Scientific, Cat# 4319413E). qPCR was performed on a CFX-Opus 384 (BioRad) using standard parameters according to the master mix instructions.

#### Image processing and analysis

Images were imported into Fiji (https://imagej.net/software/fiji/downloads, ImageJ2, version: 2.16.0/1.54p) and the size of the images adjusted to a width of 1000 pixel using “Image/Adjust/Size” and standard parameters. Subsequently, the background was subtracted using the “Subtract Background” function. Next, images (dystrophin or laminin) were imported into cellpose (https://www.cellpose.org/, version 4.0.4, platform: Darwin, python version: 3.12.9, torch version: 2.7.1) and segmented using the CPSAM algorithm. The resulting segmentation masks were visually inspected, and saved as .png files. To quantify the signal (Dystrophin or tdTomato) images were analyzed using a custom pipeline in CellProfiler (https://cellprofiler.org/, version: 4.2.6) consisting of the following modules: (i) ConvertImageToObjects (using masks as input image), (ii) MeasureObjectIntensity (measuring intensity of fibers in Dystrophin or tdTomato, respectively), and (iii) ExportToSpreadsheet (exporting all measurements into .csv files). Finally, the 99%ile of signal intensity in PBS control animals was used as a threshold to determine the percentage of fibers that surpass the threshold in the treatment conditions.

#### Lipid nanoparticle (LNP) formulation

DMG-PEG2000 (Avanti, Cat# 880151P-1g), cholesterol (Millipore Sigma, Cat# C8667-500MG), DPPC (Avanti, Cat# 850355P-25mg) were all dissolved in 100% ethanol. TCL053 (Cayman Chemical, Cat# 37045) was dissolved in methyl acetate. For muscle-optimized LNPs, a previously optimized formulation consisting of TCL053, DPPC, cholesterol, and DMG-PEG2000 (60:10.6:28.7:0.7) in ethanol was used ^21^. IVT mRNA was dissolved in 10 mM MES (2-(N-morpholino) ethanesulfonic acid) buffer (pH 5.5) at a working concentration of 0.18 mg/mL. LNPs were formulated using a NanoAssemblr® Ignite™ (Cytiva) at an aqueous-to-organic ratio of 2:1 and a flow rate of 8 ml per minute. LNPs were collected and dialyzed overnight into PBS using Slide-A-Lyzer™ Dialysis Cassettes, 10K MWCO (Thermo Fisher Scientific, Cat# 66810). LNPs were concentrated by centrifugation in a 100kD MWCO Amicon filter (Millipore Sigma, Cat# UFC810024) at 2000xg, 4C if necessary. Encapsulation efficiency was determined using a modified Ribogreen assay, encapsulation was >85% for all LNPs used in this study. Size and polydispersity of the resulting LNP were determined via dynamic light scattering on a NanoStar-II (Wyatt Technologies). LNPs had an average diameter of ∼110 nM.

