## Supplemental Information for "In vitro–reconstituted Drosophila Arc capsids deliver gene editors to dystrophic muscle"

**This PDF file includes:**
Supplementary Figure 1 – 9
Tables S1-S4

**Other Supplementary files include the following:**
Supplementary Data 1: Plasmid maps for plasmids generated in this study

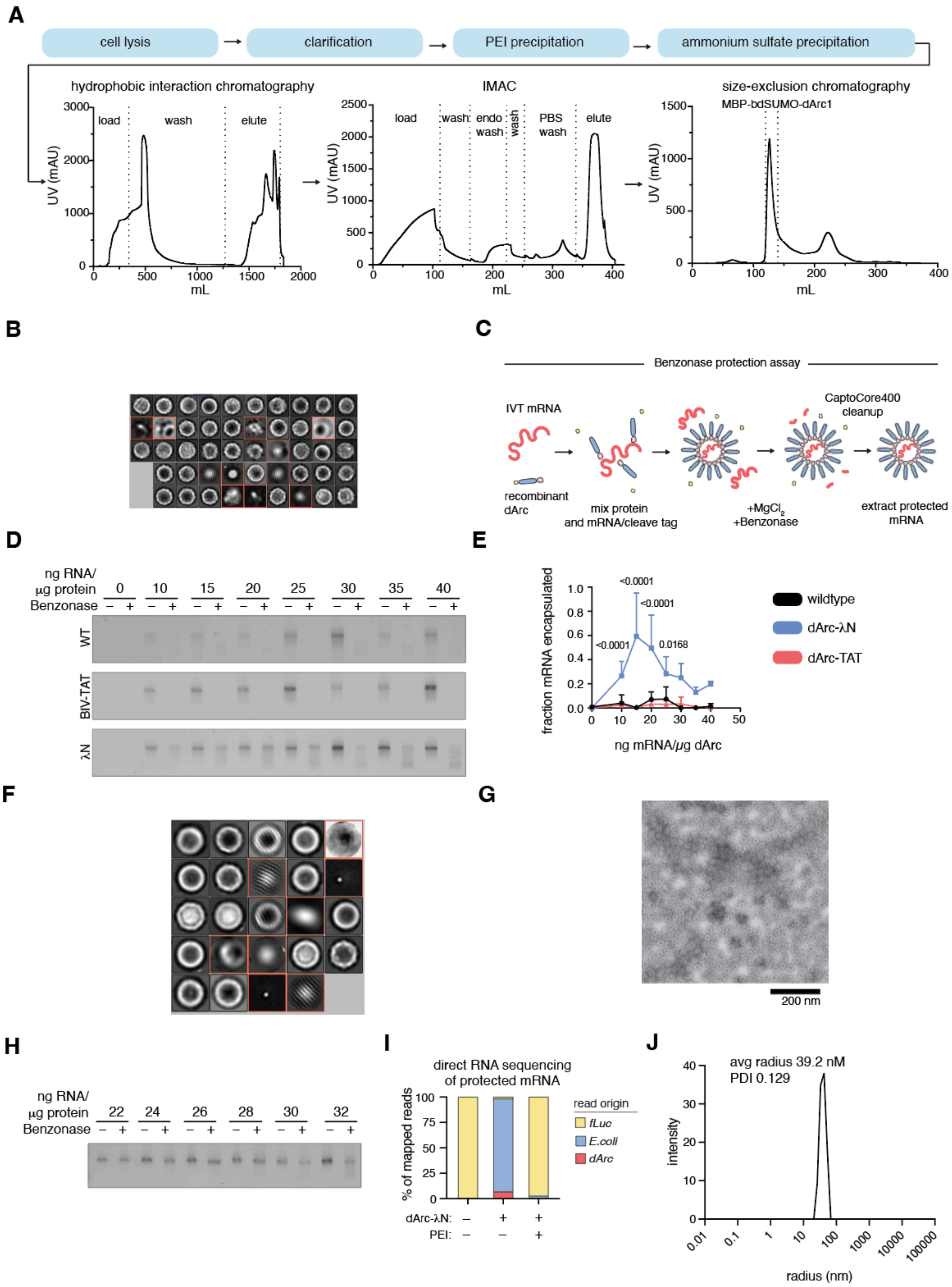
A
cell lysis
clarification
PEI precipitation
ammonium sulfate precipitation
hydrophobic interaction chromatography
load
wash
elute
UV (mAU)
mL
IMAC
load
wash
endo wash
wash
PBS wash
elute
size-exclusion chromatography
MBP-bdSUMO-dArc1
B
C
Benzonase protection assay
IVT mRNA
recombinant dArc
mix protein and mRNA/cleave tag
+MgCl2 +Benzonase
CaptoCore400 cleanup
extract protected mRNA
D
ng RNA/ µg protein
Benzonase
WT
BIV-TAT
λN
E
fraction mRNA encapsulated
ng mRNA/µg dArc
<0.0001
<0.0001
<0.0001
0.0168
wildtype
dArc-λN
dArc-TAT
F
G
200 nm
H
ng RNA/ µg protein
Benzonase
I
direct RNA sequencing of protected mRNA
% of mapped reads
read origin
fLuc
E.coli
dArc
dArc-λN:
PEI:
J
avg radius 39.2 nM
PDI 0.129
intensity
radius (nm)

**Supplementary Figure 1. dArc capsids can be engineered for efficient mRNA encapsulation**

A. Representative chromatograms from the various stages of dArc purification with PEI precipitation and endotoxin washing
B. Class averages used to quantify micrographs in Figure 1C, averages outlined in red were not used for quantification.
C. Schematic of Benzonase protection assays used to determine fraction of encapsulated mRNA by dArc capsids.
D. Representative ethidium bromide-stained agarose gels of RNA extracted from WT, λN, and BIV-TAT dArc capsids before and after Benzonase treatment at various loading ratios.
E. Quantification of ratio of encapsulated mRNA from gels in S1D. n=3 reactions per condition, two-way ANOVA.
F. Class averages used to quantify micrographs in Figure 1D, averages outlined in red were not used for quantification.
G. Electron micrograph of Spy002-dArc-λN capsids (hereafter referred to as dArc-λN). Scale bar indicates 200 nm.
H. Representative ethidium bromide-stained agarose gel of dArc-λN particles encapsulating a fLuc mRNA in the presence of 640 μM spermidine.
I. Direct RNA sequencing of RNA protected inside dArc-λN(*fLuc*$^{mRNA}$) capsids with or without PEI stripping during protein production. The first bar represents the sequencing of the input *fLuc* mRNA. fLuc, firefly Luciferase
J. Dynamic light scattering of purified dArc-λN capsids

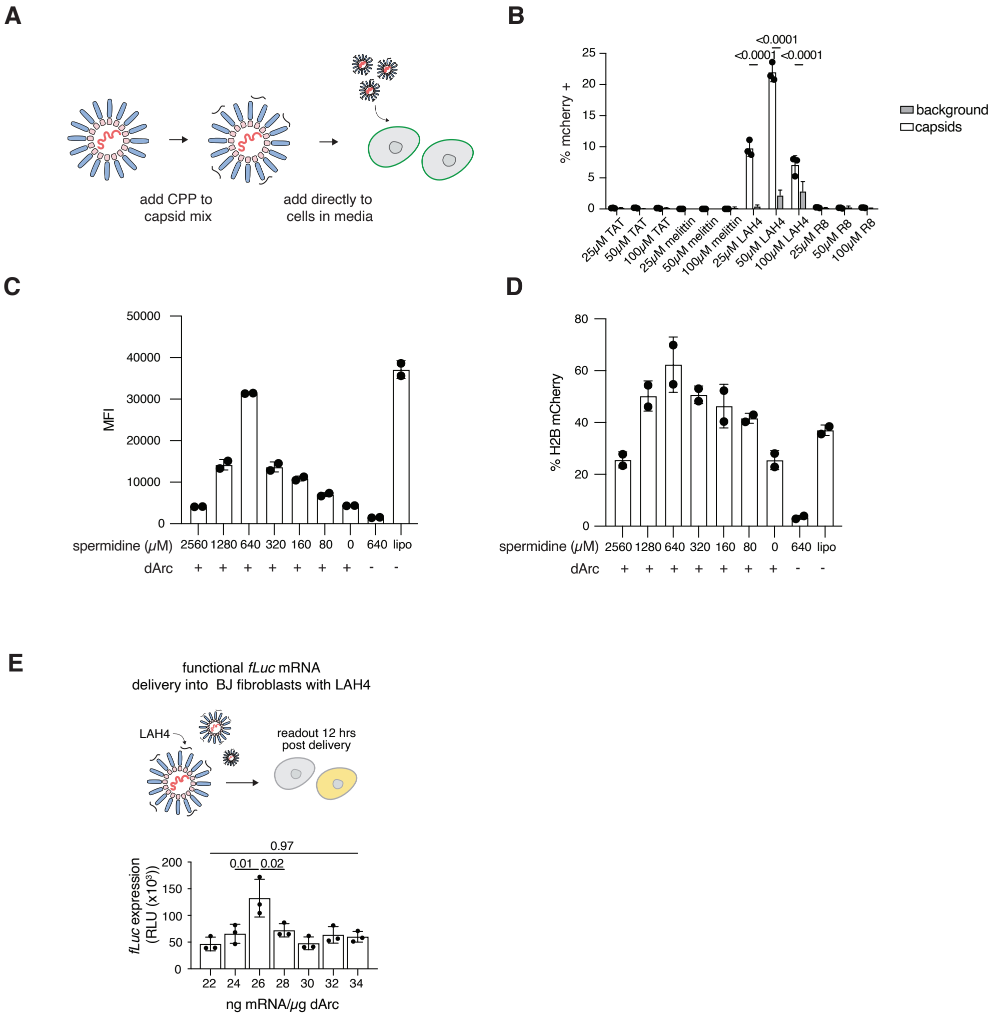

**Supplementary Figure 2. dArc can be complexed with cell-penetrating peptides to mediate functional delivery *in vitro***

A. Schematic depicting use of cell-penetrating peptides in complex with dArc to mediate functional delivery.
B. CPP-mediated delivery of dArc-λN(*H2B-mCherry*$^{mRNA}$) to BJ fibroblasts at various CPP concentrations. n=3 per condition, two-way ANOVA.
C. Mean fluorescence intensity of BJ fibroblasts treated with dArc-λN(*H2B-mCherry*$^{mRNA}$) packaged with various spermidine concentrations. Delivery mediated by LAH4.
D. Percent of H2B-mCherry positive BJ fibroblasts treated with dArc-λN(*H2B-mCherry*$^{mRNA}$) packaged with various spermidine concentrations. Delivery mediated by LAH4.
E. dArc-λN mediated delivery of protected *fLuc* mRNA into BJ fibroblasts with LAH4. Readout 16 hours post capsid treatment, one-way ANOVA with Tukey's.

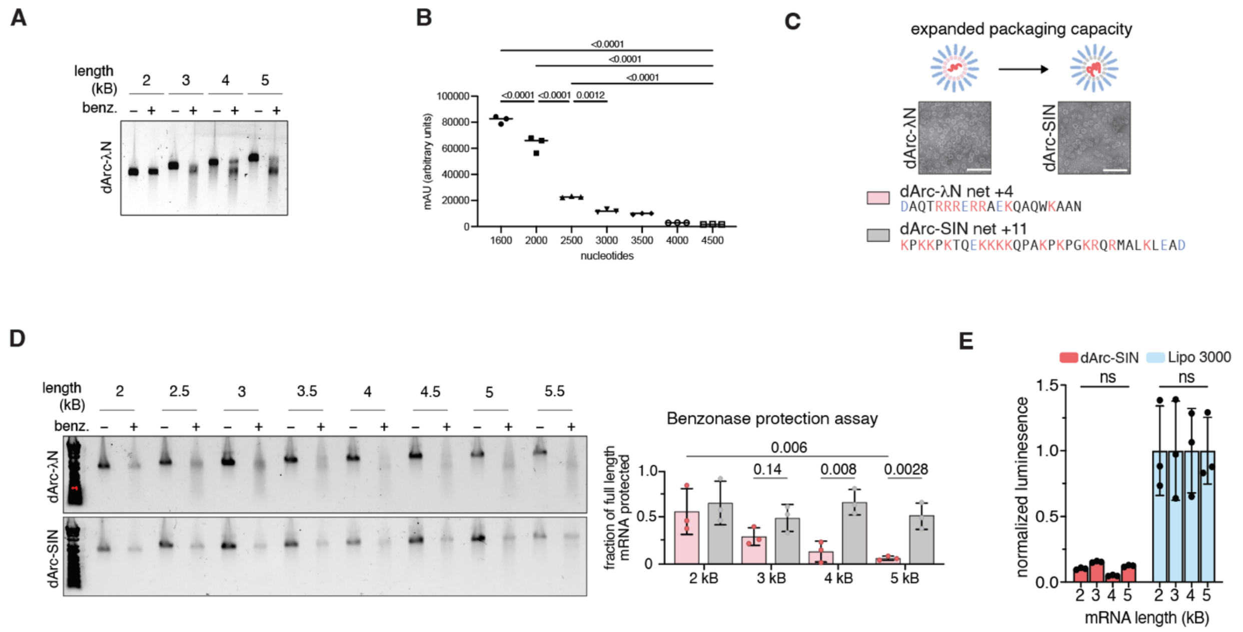

**Supplementary Figure 3. dArc can be engineered to encapsulate large mRNAs**

A. Agarose gel of RNA extracted from dArc capsids packaging various lengths of mRNA with or without benzonase treatment.
B. LAH4 mediated functional delivery of fLuc mRNAs of different length packaged in dArc-λN
C. Electron micrographs of dArc-λN and dArc-SIN capsids. Scale bar represents 200 nm.
D. (Left) Agarose gel of RNA extracted from dArc-λN and dArc-SIN capsids packaging various lengths of mRNA with or without benzonase treatment. (Right) Quantification of the fraction of protected mRNA from dArc-λN and dArc-SIN with various lengths of mRNA, n=3 per condition, two-way ANOVA with Tukey's.
E. LAH4 mediated functional delivery of fLuc mRNAs of different lengths packaged in dArc-SIN compared to transfection of the same mRNAs. n=3 per condition, one-way ANOVA with Tukey's.

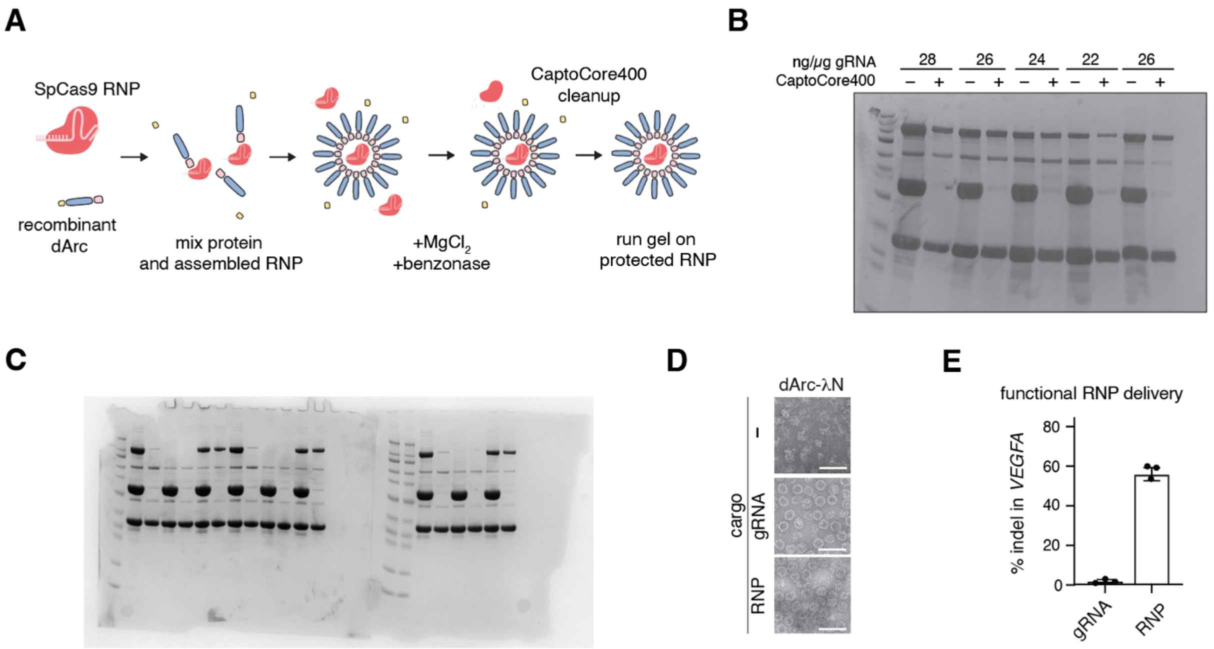

**Supplementary Figure 4. dArc can package SpCas9 RNPs**

**A.** Diagram of RNP packaging procedure
B. Coomassie blue stained SDS-PAGE gel of RNP loaded dArc-λN capsids pre or post cleanup of assembly reactions with Capto Core 400.
C. Uncropped gel from Fig. 1F
D. Electron micrographs of SpCas9 RNP complexes packaged inside dArc particles. Scale bar represents 100 nm. gRNA, dArc-λN($^{sgRNA}$); RNP, dArc-λN(Cas9$^{RNP}$)
E. Indel sequencing at *VEGFA* following delivery of SpCas9 RNPs with dArc-λN (and LAH4) to BJ fibroblasts (n=3).

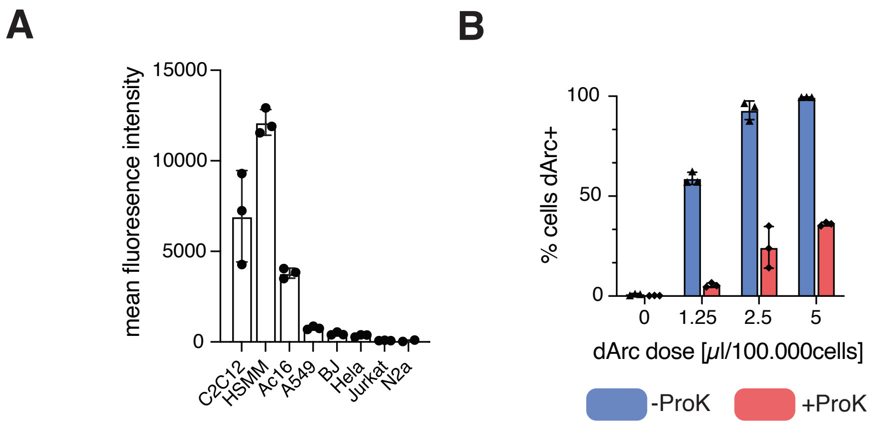

**Supplementary Figure 5. Biodistribution of dArc upon systemic injection**

A. Mean fluorescence intensity of the experiment shown in Fig. 2A
B. Percent of total cells positive for 647-dArc in C2C12 cells treated with proteinase K to remove nonbound capsids.

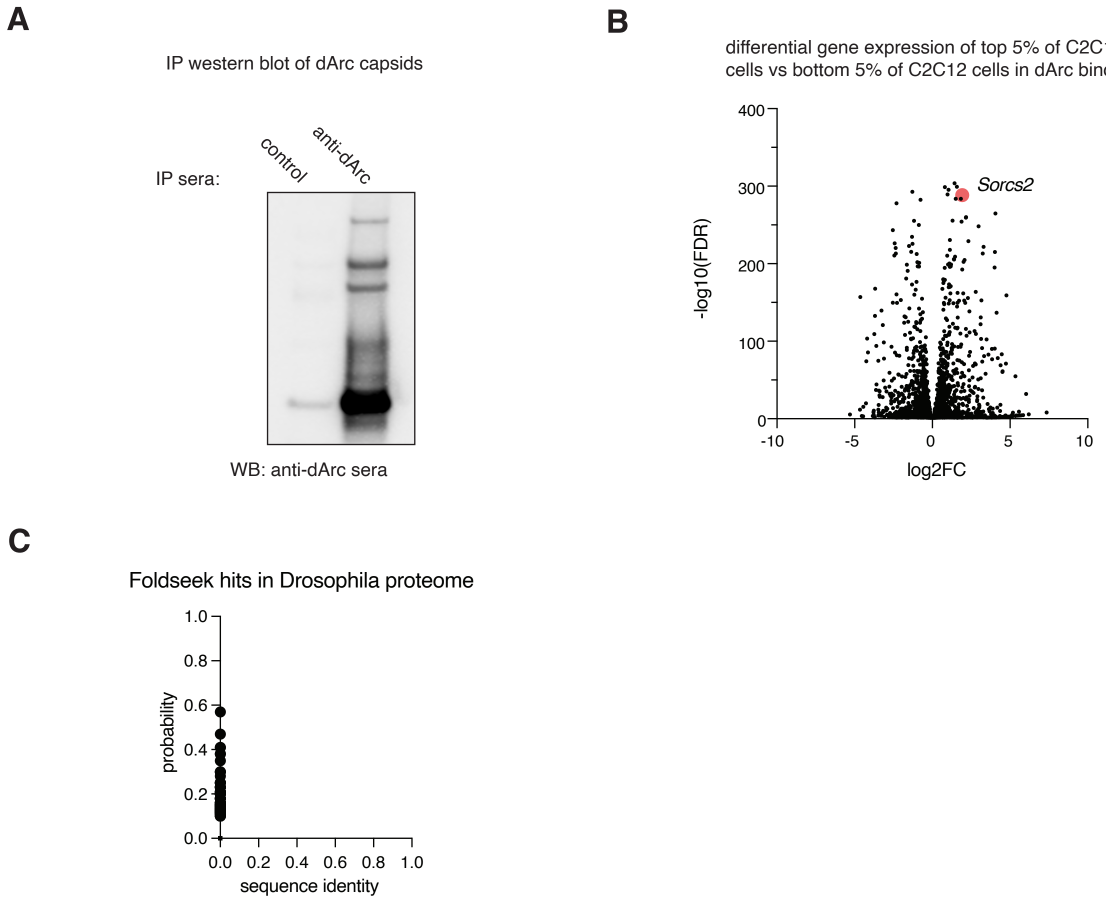

**Supplementary Figure 6. dArc directly interacts with SORCS2**

A. Co-IP western blot of dArc capsids using anti-dArc sera
B. Differential gene expression of the top 5% of cells in dArc binding vs. the bottom 5% of cells in dArc binding, Sorcs2 is denoted with a red dot and label.
C. FoldSeek results based on searching for the beta-propeller domain of mouse SORCS2 in the Drosophila proteome.

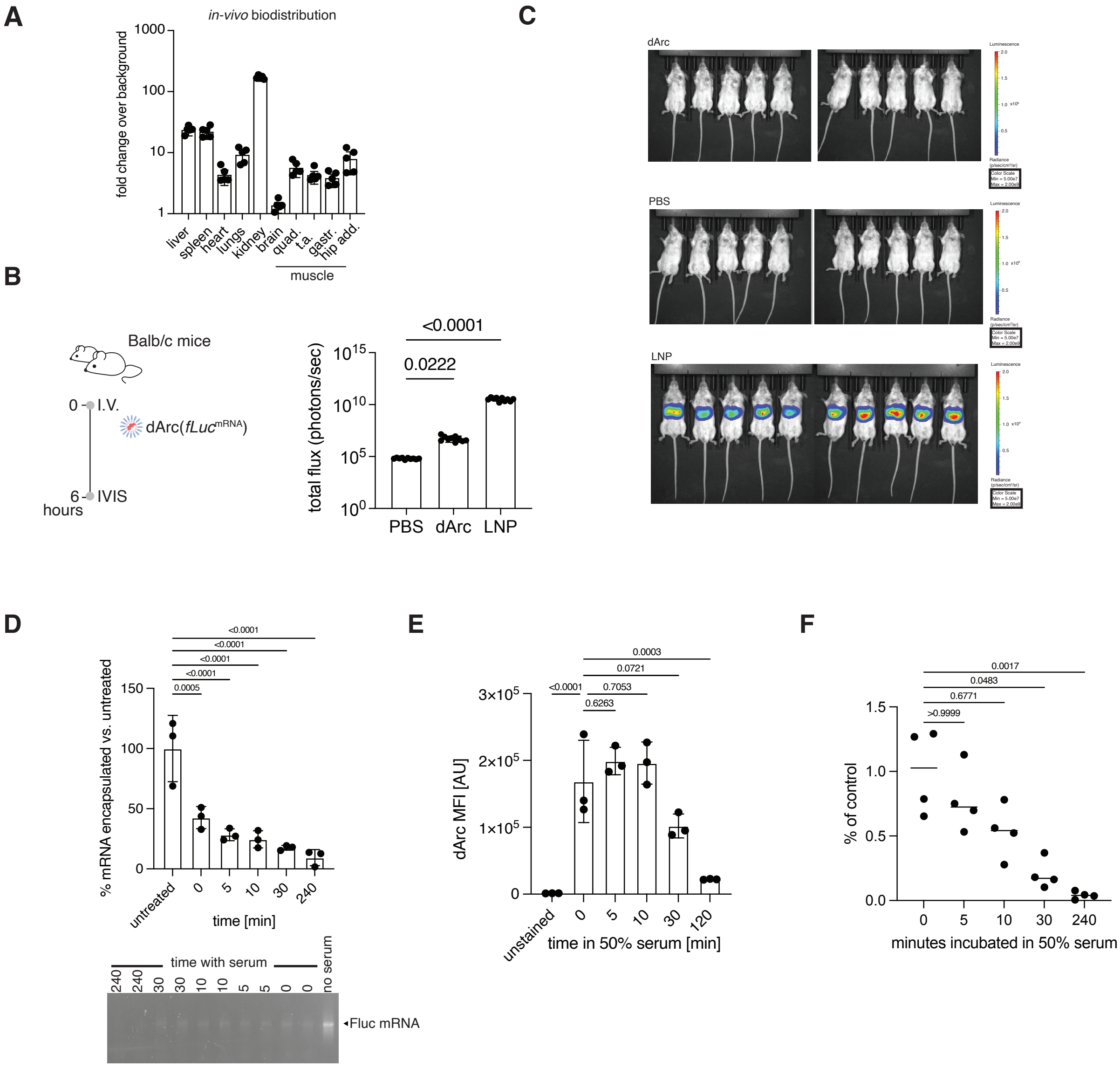

**Supplementary Figure 7. dArc particles are not efficient delivery vehicles upon systemic injection**

A. Fold change over PBS-injected animals of organs harvested from Balb/c mice injected with 0.5 mg/kg dArc(*fLuc*$^{\mathrm{IR800}}$)

B. Experimental schemiatic and quantification of liver area in images from C, Kruskal-Wallis test.

C. Bioluminesence images of Balb/c mice injected with 0.25 mg/kg dArc-λN(*fLuc*$^{\mathrm{mRNA}}$) or SM-102-fLuc LNP, scaled to LNP control.

D. (Left) Quantification of fraction of mRNA encapsulated by dArc capsids incubated in 50% mouse serum for various durations. (Right) Representative agarose gel of encapsulated mRNA. n=3 per condition, one-way ANOVA with Tukey's.

E. Mean fluorescence intensity (MFI) of dArc-λN(647-*fLuc*$^{mRNA}$) bound to C2C12 cells after pre-incubation in 50% serum for varying durations. n=3 per condition, one-way ANOVA with Tukey's.

F. Delivery of dArc-λN(*fLuc*$^{mRNA}$) to C2C12 cells after pre-incubation in 50% serum for varying durations. n=4 per condition, one-way ANOVA with Tukey's.

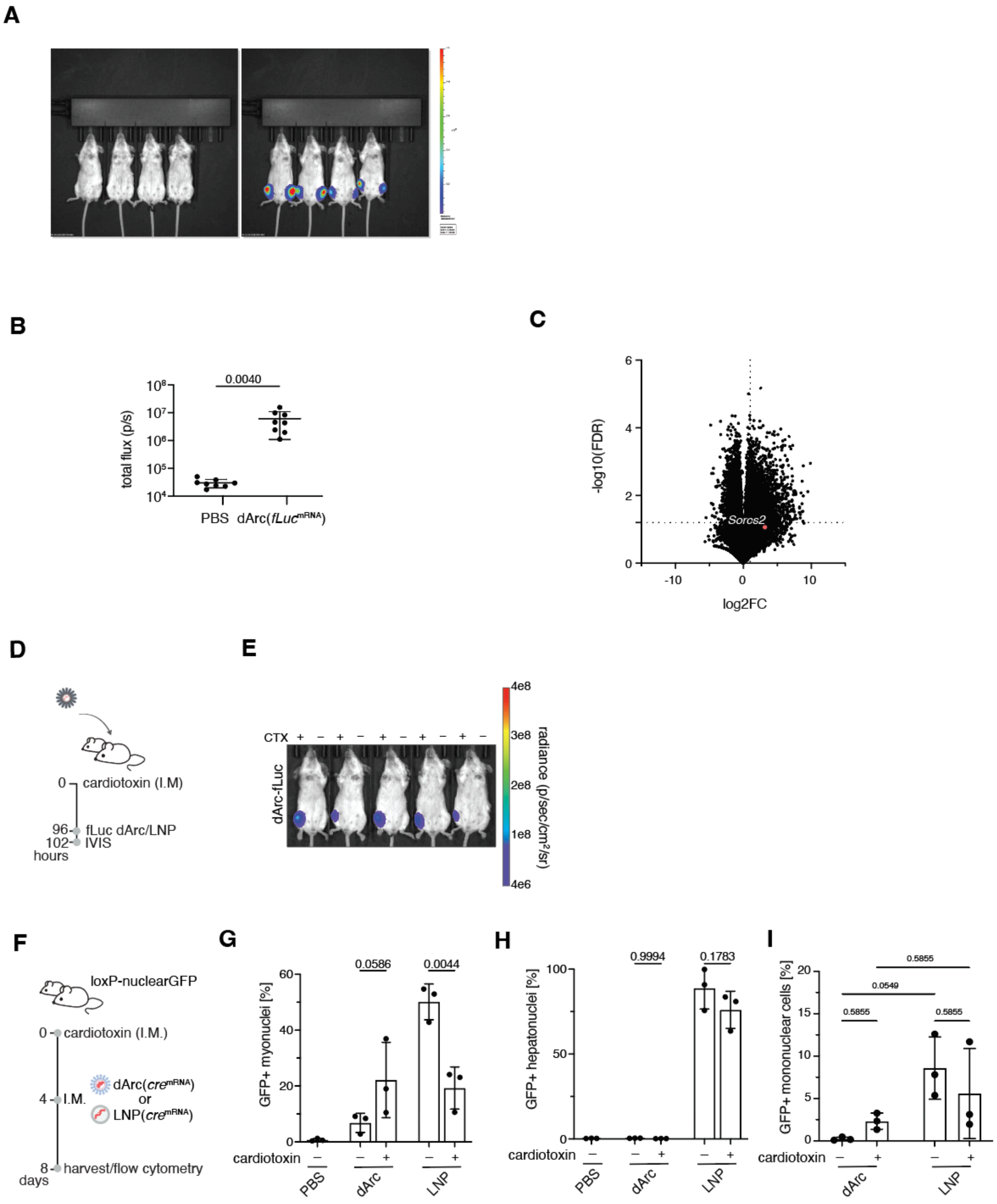

**Supplementary Figure 8. dArc can deliver to muscle *in vivo***

A. Uncropped images from Fig. 3A
B. Quantification of images from 3A
C. Differential gene expression from muscle tissue of animals at Day 4 post-cardiotoxin injury versus uninjected animals. *Sorcs2* is denoted with a red dot and label. Data and analysis are from (*20*).

D. Diagram of an experiment to investigate dArc-λN function in regenerating muscle.
E. Representative bioluminescence imaging (left) and quantification (right) of mice injected with dArc-λN(*fLuc*$^{mRNA}$) or TCL053-LNP(*fLuc*$^{mRNA}$). In each mouse, the left TA has been injured with cardiotoxin 4 days prior, whereas the right TA received a PBS injection. A subset of this data is shown in Figure 3B. Quantifications represent mean flux across n=5 animals.
F. Timeline of cardiotoxin injury model in loxP-nuclear-GFP mice
G. Quantification of PCM1+GFP+ myonuclei from mice injected I.M. with 10 μg dArc-λN(*cre*$^{mRNA}$) or 10 μg TCL053-LNP(*cre*$^{mRNA}$) four days post cardiotoxin injury, two-way ANOVA with Sidak.
H. Quantification of HNF4A+GFP+ hepatonuclei from mice injected I.M. with 10 μg dArc-λN(*cre*$^{mRNA}$) or 10 μg TCL053-LNP(*cre*$^{mRNA}$) four days post cardiotoxin injury, two-way ANOVA with Sidak.
I. Flow cytometry quantification of GFP+ mononuclear cells in muscle from mice injected with 10 ug dArc-λN(*cre*$^{mRNA}$) or 10 ug Cre- TCL-053 LNPs either in wildtype animals, or in a cardiotoxin injury model, n=3 per condition, two-way ANOVA with Sidak.

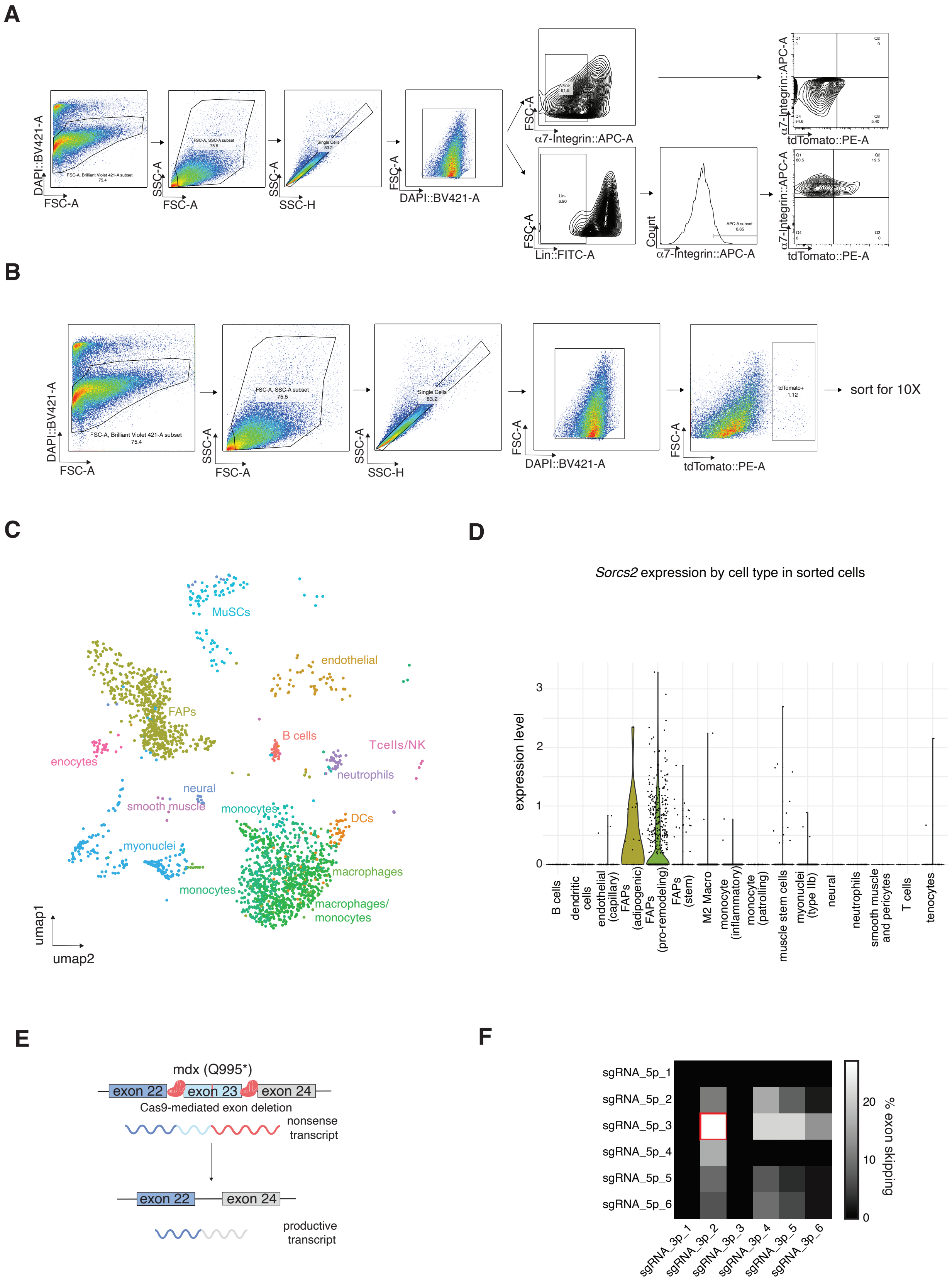
A
DAPI::BV421-A
FSC-A
SSC-A
SSC-H
DAPI::BV421-A
α7-Integrin::APC-A
tdTomato::PE-A
Lin::FITC-A
Count
B
sort for 10X
tdTomato::PE-A
C
MuSCs
endothelial
FAPs
B cells
Tcells/NK
enocytes
neutrophils
neural
smooth muscle
monocytes
DCs
myonuclei
macrophages
monocytes
macrophages/ monocytes
umap1
umap2
D
Sorcs2 expression by cell type in sorted cells
expression level
B cells
dendritic cells
endothelial (capillary)
FAPs (adipogenic)
FAPs (pro-remodeling)
FAPs (stem)
M2 Macro
monocyte (inflammatory)
monocyte (patrolling)
muscle stem cells
myonuclei (type IIb)
neural
neutrophils
smooth muscle and pericytes
T cells
tenocytes
E
mdx (Q995*)
exon 22
exon 23
exon 24
Cas9-mediated exon deletion
nonsense transcript
productive transcript
F
sgRNA_5p_1
sgRNA_5p_2
sgRNA_5p_3
sgRNA_5p_4
sgRNA_5p_5
sgRNA_5p_6
sgRNA_3p_1
sgRNA_3p_2
sgRNA_3p_3
sgRNA_3p_4
sgRNA_3p_5
sgRNA_3p_6
% exon skipping

**Supplementary Figure 9. dArc capsids can deliver to muscle progenitor cells**

A. Flow gating strategy for Fig. 3B
B. Sorting strategy for muscle mononuclear cells from Ai9 mice intramuscularly injected with dArc-λN($cre^{mRNA}$)
C. UMAP projection of single-cell RNA sequencing of muscle mononuclear cells from Ai9 mice intramuscularly injected with dArc-λN($cre^{mRNA}$) from 3 pooled animals.
D. Sorcs2 expression across clusters shown in S9C
E. Diagram of exon excision approach in mdx mice.
**F.** Exon skipping in C2C12 cells transfected with Cas9 mRNA and combinations of various exon 23 targeting five prime and three prime sgRNAs.

**Table S1** List of oligos used in this study

| Name | Sequence (5'->3') | Purpose |
|---|---|---|
| VEGFA_FWD | TCGTCGGCAGCGTCAGATGTGTATAAGA GACAGTACAGAGCTGGGTGGAGAGAGG | NGS VEGFA knockout |
| VEGFA_REV | GTCTCGTGGGCTCGGAGATGTGTATAAG AGACAGTCTTCAAGCCATCCTGTGTGC | NGS VEGFA knockout |
| DMD_FWD | TCGTCGGCAGCGTCAGATGTGTATAAG AGACAGTAGGTAAGTTAAAATGTATCA CATATATAATAAACATAGTTATTAATGC ATAGATATTCAGTA | NGS of DMD samples |
| DMD_REV | GTCTCGTGGGCTCGGAGATGTGTATAAG AGACAGCTTTGAAGGACTCTGGGTAAA ATATCTGTTTCCCA | NGS of DMD samples |
| Exon4-5_Fwd | GGCACTGCGGGTCTTACA | qPCR DMD samples |
| Exon4-5_Rev | CATCCACTATGTCAGTGCTTCCTAT | qPCR DMD samples |
| Exon22-24_Fwd | CTGAATATGAAATAATGGAGGAGAGAC TCG | qPCR DMD samples |
| Exon22-24_Rev | CTTCAGCCATCCATTTCTGTAAGGT | qPCR DMD samples |
| Exon4-5_probe | TTCACTAAATCAACATTATTTTTC | qPCR DMD samples |
| Exon22-24_probe | ATGTGATTCTGTAATTTCC | qPCR DMD samples |

**Table S2** List of plasmids used in this study (maps found in Data S1)

| Plasmid Name | Source |
|---|---|
| pMBP-bdSUMO-dArc1 | In-house generated |
| pMBP-bdSUMO-dArc1-LN | In-house generated |
| pMBP-bdSUMO-dArc1-BIVTAT | In-house generated |
| pMBP-bdSUMO-dArc1-CCMV | In-house generated |
| pMBP-bdSUMO-dArc1-delZF | In-house generated |
| pMBP-bdSUMO-dArc1-LN11-36 | In-house generated |
| pMBP-bdSUMO-Spy002-dArc-LN | In-house generated |
| pMBP-bdSUMO-Spy002-dArc-SIN | In-house generated |
| pCMV-dArc-3'UTR | In-house generated |
| pIVT-fLuc-BXB | In-house generated |
| pIVT-H2B-mCherry-BXB | In-house generated |
| pIVT-Cre-BXB | In-house generated |
| pIVT-fLuc-SIN | In-house generated |
| pIVT-SpCas9-SIN | In-house generated |
| psPAX2 | Addgene plasmid # 12260 |
| pMD2.G | Addgene plasmid # 12259 |
| pLVX-mSorcs2-IRES-Puro | In-house generated |
| pCMV-mSorcs2ecto-Fc | In-house generated |

| | |
|---|---|
| pCMV-mSorcs2ecto_deltaNterm-Fc | In-house generated |
| pCMV-mSorcs2ecto_deltaVPS10-Fc | In-house generated |
| pCMV-mSorcs2ecto_deltaPKDN-Fc | In-house generated |
| pCMV-mSorcs2ecto_deltaPKDC-Fc | In-house generated |
| pCMV-mSorcs2ecto_deltaC-Fc | In-house generated |
| pCMV-Fc | In-house generated |

**Table S3 dArc protein sequences used in this study**

| **Name** | **Amino Acid Sequence** | **Notes** |
|---|---|---|
| dArc wildtype | AQLTQMTNEQLRELIEAVRAAAVGAA<br>GSAAAAGGADASRGKGNFSACTHSFG<br>GTRDHDVVEEFIGNIETYKDVEGISDE<br>NALKGISLLFYGMASTWWQGVRKEAT<br>TWKEAIALIREHFSPTKPAYQIYMEFFQ<br>NKQDDHDPIDTFVIQKRALLAQLPSGR<br>HDEETELDLLFGLLNIKYRKHISRHSV<br>HTFKDLLEQGRIIEHNNQEDEEQLATA<br>KNTRGSKRTTRCTYCSFRGHTFDNCR<br>KRQKDRQEEQHEE | |
| dArc-lambda N | AQLTQMTNEQLRELIEAVRAAAVGAA<br>GSAAAAGGADASRGKGNFSACTHSFG<br>GTRDHDVVEEFIGNIETYKDVEGISDE<br>NALKGISLLFYGMASTWWQGVRKEAT<br>TWKEAIALIREHFSPTKPAYQIYMEFFQ<br>NKQDDHDPIDTFVIQKRALLAQLPSGR<br>HDEETELDLLFGLLNIKYRKHISRHSV<br>HTFKDLLEQGRIIEHNNQEDEEQLATA<br>KNTRGSKRTTDAQTRRRERRAEKQAQ<br>WKAAN | Corresponds to residues 2-22 of lambda phage Antitermination protein N |
| dArc-CCMV | AQLTQMTNEQLRELIEAVRAAAVGAA<br>GSAAAAGGADASRGKGNFSACTHSFG<br>GTRDHDVVEEFIGNIETYKDVEGISDE<br>NALKGISLLFYGMASTWWQGVRKEAT<br>TWKEAIALIREHFSPTKPAYQIYMEFFQ<br>NKQDDHDPIDTFVIQKRALLAQLPSGR | |

| | | |
|---|---|---|
| | HDEETELDLLFGLLNIKYRKHISRHSV<br>HTFKDLLEQGRIIEHNNQEDEEQLATA<br>KNTRGSKRTT<u>RAQRRAAARKNKRNTR</u><br><u>VVQP</u> | |
| dArc-BIV TAT | AQLTQMTNEQLRELIEAVRAAAVGAA<br>GSAAAAGGADASRGKGNFSACTHSFG<br>GTRDHDVVEEFIGNIETYKDVEGISDE<br>NALKGISLLFYGMASTWWQGVRKEAT<br>TWKEAIALIREHFSPTKPAYQIYMEFFQ<br>NKQDDHDPIDTFVIQKRALLAQLPSGR<br>HDEETELDLLFGLLNIKYRKHISRHSV<br>HTFKDLLEQGRIIEHNNQEDEEQLATA<br>KNTRGSKRTT<u>SGPRPRGTRGKGRRIRR</u> | |
| dArc-delZF | AQLTQMTNEQLRELIEAVRAAAVGAA<br>GSAAAAGGADASRGKGNFSACTHSFG<br>GTRDHDVVEEFIGNIETYKDVEGISDE<br>NALKGISLLFYGMASTWWQGVRKEAT<br>TWKEAIALIREHFSPTKPAYQIYMEFFQ<br>NKQDDHDPIDTFVIQKRALLAQLPSGR<br>HDEETELDLLFGLLNIKYRKHISRHSV<br>HTFKDLLEQGRIIEHNNQEDEEQLATA<br>KNTRGSKRTT | |
| dArc-LN11-36 | AQLTQMTNEQLRELIEAVRAAAVGAA<br>GSAAAAGGADASRGKGNFSACTHSFG<br>GTRDHDVVEEFIGNIETYKDVEGISDE<br>NALKGISLLFYGMASTWWQGVRKEAT<br>TWKEAIALIREHFSPTKPAYQIYMEFFQ<br>NKQDDHDPIDTFVIQKRALLAQLPSGR<br>HDEETELDLLFGLLNIKYRKHISRHSV<br>HTFKDLLEQGRIIEHNNQEDEEQLATA<br>KNTRGSKRTT<u>RAEKQAQWKAANPLL</u><br><u>VGVSAKPVNRP</u> | |
| Spy002-dArc-lambda N | VPTIVMVDAYKRYK*SGSETPGTSESATP*<br>*ES*AQLTQMTNEQLRELIEAVRAAAVG<br>AAGSAAAAGGADASRGKGNFSACTHS<br>FGGTRDHDVVEEFIGNIETYKDVEGIS<br>DENALKGISLLFYGMASTWWQGVRKE<br>ATTWKEAIALIREHFSPTKPAYQIYMEF<br>FQNKQDDHDPIDTFVIQKRALLAQLPS<br>GRHDEETELDLLFGLLNIKYRKHISRHS<br>VHTFKDLLEQGRIIEHNNQEDEEQLAT<br>AKNTRGSKRTT<u>DAQTRRRERRAEKQA</u> | used for all lambda N figures after 2D |

| | |
|---|---|
| | QWKAAN |
| Spy002-dArc-SIN | VPTIVMVDAYKRYK*SGSETPGTSESATPES*AQLTQMTNEQLRELIEAVRAAAVGAAGSAAAAGGADASRGKGNFSACTHSFGGTRDHDVVEEFIGNIETYKDVEGISDENALKGISLLFYGMASTWWQGVRKEATTWKEAIALIREHFSPTKPAYQIYMEFFQNKQDDHDPIDTFVIQKRALLAQLPSGRHDEETELDLLFGLLNIKYRKHISRHSVHTFKDLLEQGRIIEHNNQEDEEQLATAKNTRGSKRTTKPKKPKTQEKKKKQPAKPKPGKRQRMALKLEAD* |

**Table S4 Guide RNAs used in this study**

| Name | Sequence (5'->3') | Species |
|---|---|---|
| VEGFA | TCATGCAGTGGTGAAGTTCA | Human |
| DMD5p_1 | TTAAGCTTAGGTAAAATCAA | Mouse |
| DMD5p_2 | TTATTTTAATAGCCTAAGTC | Mouse |
| DMD5p_3 (used in vivo) | TCTTAATAATGTTTCACTGT | Mouse |
| DMD5p_4 | TTTCATTCATATCAAGAAGA | Mouse |
| DMD5p_5 | AATAATTTCTATTATATTAC | Mouse |
| DMD5p_6 | ATAATTTCTATTATATTACA | Mouse |
| DMD3p_1 | GATCATGGATTTGACACTTT | Mouse |
| DMD3p_2 (used in vivo) | ATGTTAAGTATACTTGGAGT | Mouse |
| DMD3p_3 | GAATGATCAAGTCACTAGCA | Mouse |
| DMD3p_4 | CGAAAATTTCAGGTAAGCCG | Mouse |
| DMD3p_5 | ATAGTTTAAAGGCCAAACCT | Mouse |
| DMD3p_6 | TTTTTCACATAGCAATTAAT | Mouse |
